# AMPK senses cellular levels of nicotinamide adenine dinucleotide

**DOI:** 10.64898/2026.05.19.726234

**Authors:** Niall Wilson, Yoana Rabanal Ruiz, Aniketh Bishnu, Shangze Xu, Congxin Sun, Mohd Syed Ahangar, Kevin M. Rattigan, Jiangyu Tang, Danyelle Silva-Amaral, Conchita Fraguas Bringas, Kei Sakamoto, Tetsushi Kataura, Sovan Sarkar, Elton Zeqiraj, G. Vignir Helgason, Ian G. Ganley, Agnieszka K. Bronowska, Viktor I. Korolchuk

## Abstract

The electron shuttle and coenzyme nicotinamide adenine nucleotide (NAD) is essential for cellular metabolism and homeostasis. NAD levels significantly fluctuate in cells, whilst several age-related diseases are associated with depletion of this metabolite. However, how NAD changes are monitored by nutrient/energy sensing signalling pathways remains poorly understood. We found that at physiological concentrations NAD controls the activity of the AMP-activated protein kinase (AMPK) *in vitro* and in human cells. Mechanistically, NAD binds gamma subunit of AMPK, and mutagenesis of the putative binding site renders the holoenzyme insensitive to NAD inhibition. Hyperactivation of AMPK in response to NAD depletion suppresses metabolic pathways including mammalian Target of Rapamycin Complex I (mTORC1) and autophagy. These results demonstrate that in addition to monitoring cellular energy levels AMPK functions as a NAD sensor, providing novel insight into how cells and tissues detect and respond to metabolic fluctuations with implications for stress resistance and ageing.

## Introduction

NAD is an essential metabolic intermediate and enzyme cofactor central to energy homeostasis. Both total intracellular NAD concentrations and redox status fluctuate in response to nutrient availability, metabolic activity, circadian rhythms, and stress conditions [1–4]. Cellular levels of the NAD pool have also been shown to decline in age-related neurodegenerative and metabolic diseases [5–7]. Beyond its classical role in electron transfer reactions, the oxidised form of NAD (NAD⁺) functions as a critical signalling metabolite whose availability directly determines the activity of several NAD⁺-consuming enzymes. These include sirtuins (e.g. SIRT1), poly(ADP-ribose) polymerases (PARPs), and NADases such as CD38, which compete for a common NAD⁺ pool and couple metabolic state to chromatin remodelling, DNA repair, stress adaptation, and inflammatory signalling [3,7,8].

Despite these established NAD^+^-consuming pathways, how cells sense changes in NAD availability and coordinate NAD homeostasis with broader metabolic signalling remains incompletely understood. Cellular NAD⁺ levels are maintained through a dynamic balance between biosynthesis, recycling and consumption. The predominant salvage pathway, controlled by the rate-limiting enzyme nicotinamide phosphoribosyltransferase (NAMPT), replenishes NAD⁺ from nicotinamide and is regulated by nutrient availability and stress-response pathways [9,10]. AMP-activated protein kinase (AMPK), a serine/threonine kinase that senses cellular energy status through changes in AMP, ADP and ATP, can promote NAD⁺ synthesis by increasing NAMPT expression and directly phosphorylating NAMPT to increase its activity [11,12]. Beyond its role in NAD^+^ metabolism, AMPK also regulates mechanistic Target of Rapamycin Complex 1 (mTORC1) signalling and macroautophagy (hereinafter autophagy), a lysosome-dependent degradative pathway essential for cellular quality control [13,14]. Although AMPK has historically been characterised as an activator of autophagy under energy stress, recent data indicate a more complex relationship, with AMPK reported to both activate and inhibit autophagy depending on the stimulus [15–18].

In addition to classical adenylates, sensing of other nucleotides including NAD by AMPK has been investigated in the past. NAD was shown to bind AMPK at the γ-subunit cystathionine β-synthase (CBS) sites, however conflicting data was reported regarding its impact on AMPK catalytic activity [19–22]. As such, whether cellular NAD availability acts as a physiologically relevant input into AMPK signalling, and whether this regulates cellular metabolic status, remains unresolved. Here we show that AMPK signalling is responsive to cellular NAD deficit, driving its activation independently of detectable changes in adenylate energy status. Mechanistically, NAD⁺ suppresses AMPK activity *in vitro* and in cells, and mutation within the γ1 regulatory subunit renders the holoenzyme insensitive to NAD⁺ inhibition. Hyperactivation of AMPK in response to NAD depletion shuts down both anabolic (mTORC1) and catabolic (autophagy) pathways. These findings identify AMPK as a direct NAD sensor and provide mechanistic insight into how NAD depletion associated with stress and age-related disease negatively impacts metabolic activity.

## Results

### NAD depletion suppresses autophagy initiation

Previous studies have suggested that the cellular response to NAD⁺ depletion is primarily mediated by NAD⁺-dependent proteins such as SIRT1 and PARP1. Activity of these enzymes declines under NAD⁺ deficit, leading to suppression of metabolic pathways including autophagy [3]. To model NAD depletion we used FK866 as a selective inhibitor of NAMPT, a rate-limiting enzyme in the NAD^+^ salvage pathway. FK866 alone and in combination with nicotinamide riboside (NR) served to specifically identify NAD-dependent responses (Fig. 1A). FK866 treatment in human ARPE-19 cells strongly depleted total levels of the NAD (NAD^+^/NADH) pool, with this effect rescued by NR cotreatment (Fig. 1B). Halo-GFP-LC3 reporter was used to measure autophagy responses to NAD perturbation under basal conditions and following rapamycin-induced autophagy stimulation (Fig. 1C). Lysosomal cleavage of Halo-GFP-LC3 into Halo monomer assessed by immunoblotting, and Halo^+^/GFP^-^ puncta visualised by fluorescence microscopy indicated significant suppression of both basal and rapamycin-stimulated autophagy activity in response to FK866 treatment (Fig. 1D-G). An endogenous LC3-II flux assay confirmed the autophagy deficit (Fig. 1H, I). Similar results were found in an alternative human HMC3 cell line (Fig. S1A-E).

**Figure 1.**
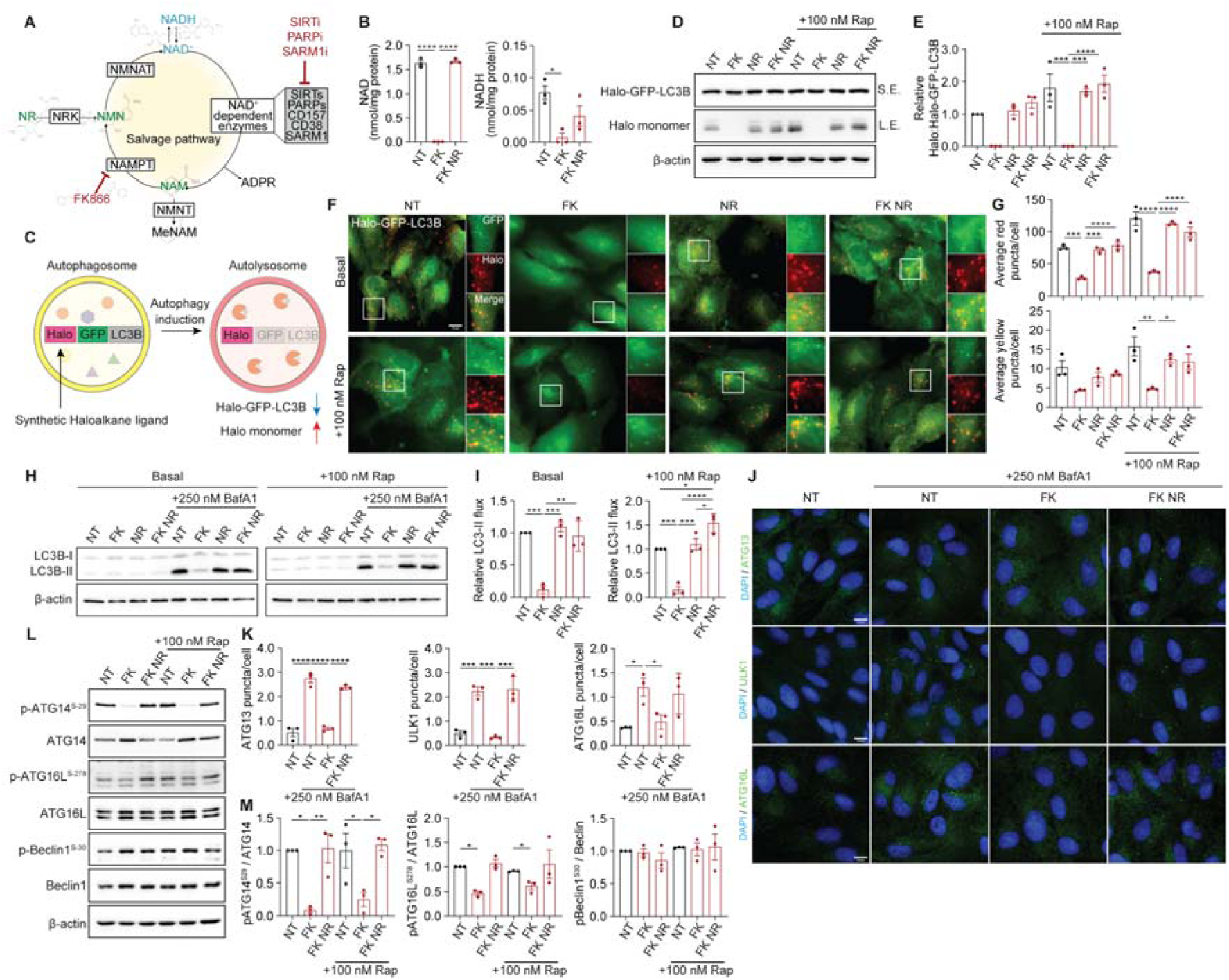
FK866-induced NAD depletion blocks autophagosome formation. **A,** Graphical representation of the NAD salvage pathway. NAD precursor molecules are highlighted in green, and enzyme inhibitors in red. **B,** Measurement of NAD^+^ and NADH in ARPE-19 cells treated with 100 nM FK866 (FK) in the presence or absence of 1 mM nicotinamide riboside (NR) for 72 h. **C,** Schematic illustration of the Halo processing assay. **D, E,** Representative immunoblot (D) and quantification (E) of Halo processing assay in ARPE-19 cells stably expressing Halo-GFP-LC3B, treated with 100 nM FK and/or 1 mM NR for 72 h, with 100 nM rapamycin (Rap) added for the final 6 h. **F, G**, Fluorescence microscopy images (F) and quantification (G) of autophagy events (red) and autophagosomes (yellow) in ARPE-19 cells stably expressing Halo-GFP-LC3B, treated as in D. **H, I,** Immunoblot analysis (H) and quantification (I) of autophagy flux in ARPE-19 cells treated as in C. **J, K,** Fluorescence microscopy images (J) and puncta quantification (K) of fixed ARPE-19 cells labelled with an anti-ATG13, anti-ULK1 or anti-ATG16L antibody. Cells were treated with as in F, with bafilomycin A1 (BafA1) added for the final 3 h of treatment to increase baseline puncta number. **L, M,** Representative immunoblots (L) and quantifications (M) of indicated proteins from ARPE-19 cells treated as indicated for 72 h, with 100 nM Rap added for the final 6 h. *P* values were calculated by one-way ANOVA and Tukey’s multiple comparisons test, on three independent experiments. \**P*<0.05; \*\**P*<0.01; \*\*\**P*<0.001; \*\*\*\**P*<0.0001. Scale bars: 10 µm (**F, J**). See also Figure S1.

Autophagy impairment in response to FK866 treatment occurred at the initiation stage mediated by the intracellular ULK1 complex and ATG16 puncta formation (Fig. 1J, K, Fig. S1F). Phosphorylation of ATG16 and ATG14, downstream targets of ULK1 complex, was also suppressed (Fig. 1L, M, Fig. S1G). Thus, FK866 treatment and the subsequent NAD loss results in autophagy impairment, rescuable by NR supplementation. However, inhibition of NAD^+^-dependent enzymes SIRT1 and PARP1, separately or in combination, did not phenocopy the effect of NAD depletion on autophagy (Fig. S1H-J).

### AMPK senses intracellular NAD levels

Given that NAD depletion impaired autophagy induction, we next measured the activity of nutrient/energy sensing pathways converging on the ULK1 complex. Depletion of NAD pools by pharmacological or genetic inhibition of NAMPT increased AMPK and suppressed mTORC1 pathway activity indicating a starvation-like response both in ARPE-19 and HMC3 cells (Fig. S2A-C). To gain insight into the kinetics of cellular response to NAD deficit, we performed analyses of metabolites and signalling pathways in ARPE-19 cells at three time points following FK866 treatment (Fig. 2A, B). Metabolomics confirmed rapid (24 hours) and persistent depletion of NAD pools and a complete rescue of this effect by NR (Fig. 2A). At the same time ATP depletion and the concomitant elevated AMP/ATP ratio, an established signal for AMPK activation, did not occur until 48 hours after the treatment (Fig. 2A, Fig. S2D). Perturbation of other metabolites, including methyl nicotinamide, glycolytic intermediates and amino acids, also became apparent at later time points (Fig. S2D-F).

**Figure 2.**
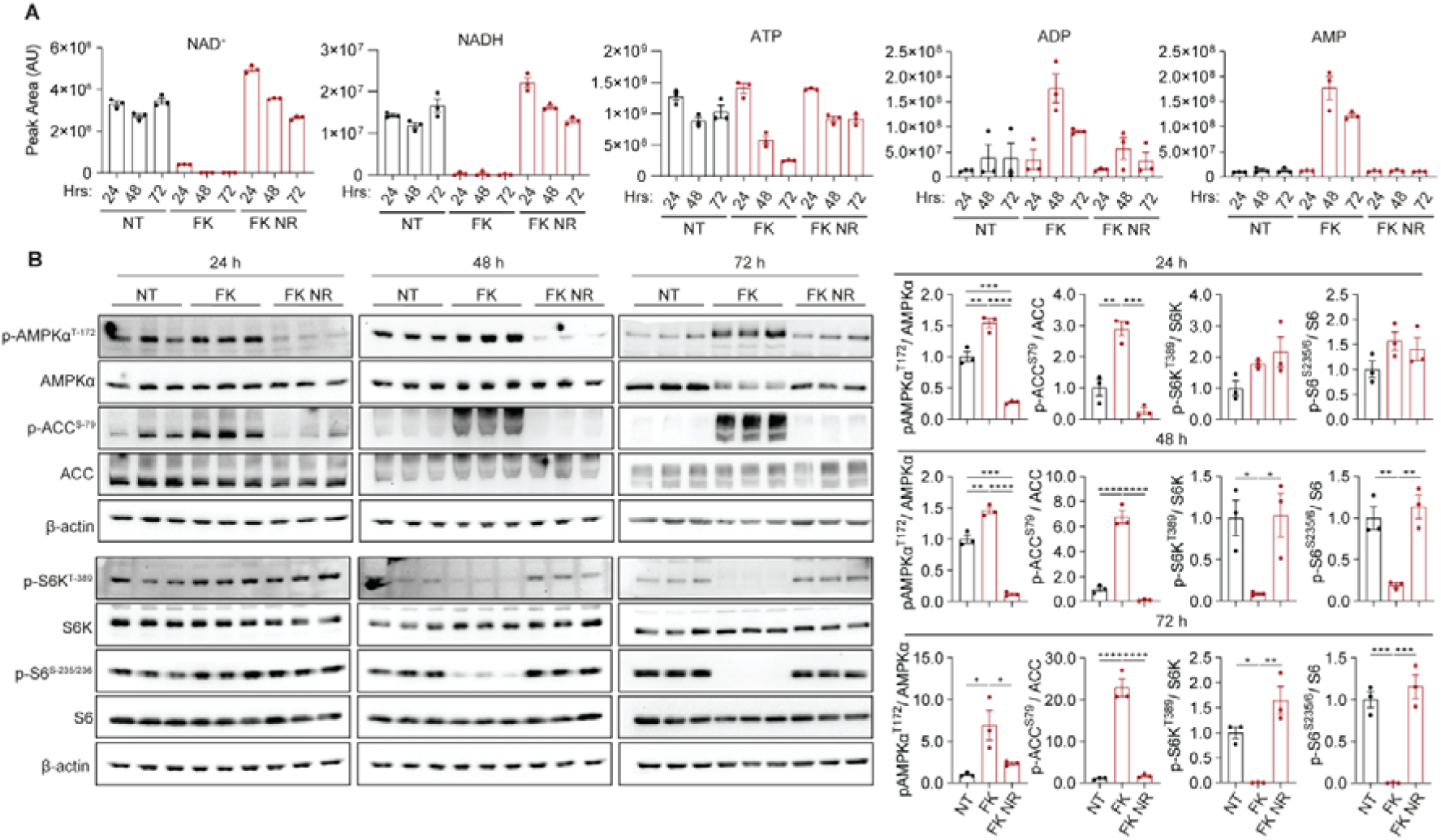
NAD depletion activates AMPK independently of changes to the AMP/ATP ratio. **A,** Metabolite analysis in ARPE-19 cells by LC-MS. Cells were treated with 100 nM FK866 (FK) in the presence of absence of 1 mM nicotinamide riboside (NR) for the indicated timepoints. Lysates were normalised to 2 × 10L cells/mL. **B,** Immunoblots and quantifications of indicated AMPK and mTOR-related proteins from ARPE-19 cells treated as in A for 24, 48 and 72 h. *P* values were calculated by Ordinary one-way ANOVA and Tukey’s multiple comparisons test. \**P*<0.05; \*\**P*<0.01; \*\*\**P*<0.001; \*\*\*\**P*<0.0001. All immunoblots in this figure represent three independent biological replicates per treatment condition (n = 3 per condition, 9 samples total).

Analyses of signalling responses determined that AMPK activation occurs 24 hours after FK866 treatment, which coincides with NAD^+^/NADH depletion in the absence of detectable perturbation of other metabolites (Fig. 2A, B, Fig. S2D-F). Thus, in our experimental setup AMPK is activated when cellular NAD but not ATP pools become depleted. In contrast, mTORC1 activity declined at the later (48 hour) timepoint, and this effect occurred in the absence of response by the upstream AKT signalling (Fig. 2B, Fig. S2G). Consistent with this, FK866 reduced phosphorylation of ULK1 at the mTORC1 site Ser757, whereas the effect on the AMPK site Ser556, although not reaching significance, was suppressed by NR co-treatment (Fig. S2H). These data suggest that NAD depletion engages AMPK while subsequently suppressing mTORC1-dependent signalling to ULK1. To test whether mTORC1 suppression was AMPK-dependent, we used cells with the knockout of α1 and α2 AMPK subunits (DKO, Fig. S2I). FK866 treatment did not suppress mTORC1 activity in the absence of AMPK (Fig. S2J). Together, these findings identify AMPK as a sensor of cellular NAD.

### AMPK activity is suppressed by NAD^+^

Previous reports provided conflicting data on the regulation of AMPK activity by NAD [19,22], and its physiological relevance, if any, is unknown. To investigate the effect of NAD on AMPK activity *in vitro*, we used luminescence-based ADP-Glo assay measuring phosphorylation of SAMS (HMRSAMSGLHLVKRR) peptide by the recombinant AMPK holoenzyme (Fig. S3A). We observed dose-dependent inhibition of AMPK activity by NAD^+^ in the presence of 100 μM AMP, with inhibition of AMPK by BAY-3827 in the same conditions serving as a positive control (Fig. 3A, B). The IC50 for NAD^+^ (∼300 μM) was found to be within the physiological range of intracellular NAD^+^ (100-1,000 μM) indicating possibility of a direct interaction with this metabolite by AMPK *in vivo* [23]. Additional assay controls confirmed specificity of the AMPK-dependent effect of NAD^+^ on the luminescence readout both in the presence and absence of AMP and AMPK (Fig. S3B, C).

**Figure 3.**
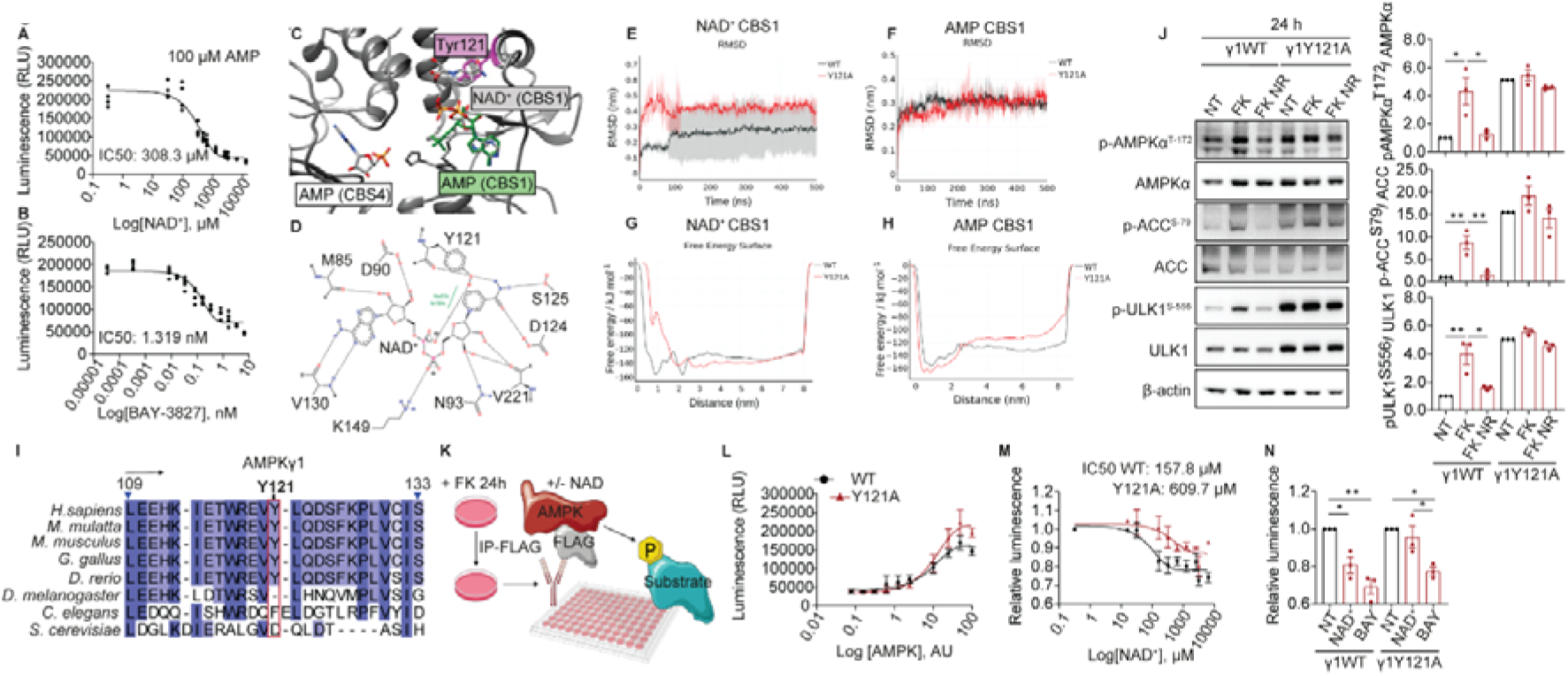
NAD^+^ suppresses AMPK activity. A-B, Kinase activity of recombinant human AMPKα1β1γ1 measured using the ADP-Glo™ assay in the presence of 100 μM AMP. AMPK (10 ng) was pre-incubated for 1 h with NAD^+^ (0-13,000 μM) (A) or BAY-3827 (0-10 nM) (B) prior to initiating the kinase reaction. Data represent three technical replicates. **C,** Predicted NAD^+^ binding mode at AMPKγ1 CBS1 site. NAD^+^ (coloured by element) is superimposed with AMP (green) at CBS1 to show overlap of their AMP moieties. Tyr121, predicted to interact with the NAD^+^ nicotinamide moiety, is shown in purple; AMPKγ1 backbone in grey. **D,** 2D interaction diagram of NAD^+^ bound at CBS1. Residues involved in stabilising interactions are labelled. Hydrophobic interactions are coloured in green, hydrogen bonds and salt bridges are dashed lines. Pi-pi interactions are omitted. **E-F,** Average root-mean-square deviation (RMSD) of NAD^+^ (E) or AMP (F) bound at CBS1 in wild-type (WT, black) and Y121A mutant (red) during 500 ns equilibrium all-atom MD simulations. Data represent averages of three independent replicas; shaded areas are SD. **G-H,** Reconstructed free energy surfaces for simulated unbinding of NAD^+^ (G) or AMP (H) from CBS1 of WT (black) and Y121A mutant (red) AMPK, calculated by well-tempered metadynamics. Y121A selectively altered the free energy profile of NAD^+^ binding, while AMP binding remained largely unchanged. **I,** Alignment showing tyrosine residue 121 (highlighted red) in AMPKγ1 across selected eukaryotes. Increasing conservation across species is shown by light-dark blue. **J,** Representative immunoblots and quantifications of indicated proteins from ARPE-19 cells stably expressing WT or Y121A mutant FLAG-AMPKγ1, treated with 100 nM FK866 (FK) in the presence or absence of 1 mM nicotinamide riboside (NR) for 24 h. The NT baseline of Y121A was scaled to the average NT of WT to account of its higher activity. **K,** Schematic of the experimental procedure to measure the kinase activity of immunoprecipitated AMPK complexes containing WT or Y121A mutant FLAG-AMPKγ1. **L,** Kinase activity of immunoprecipitated AMPK containing the WT or Y121A mutant FLAG-AMPKγ1 subunit measured using the ADP-Glo™ assay in the presence of 100 μM AMP. The X-axis represents log[AMPK] in arbitrary units, as the enzyme was immobilised on beads. Cells were treated with 100 nM FK for 24 h prior to lysis and immunoprecipitation. **M,** Kinase activity of immunoprecipitated AMPK containing WT or Y121A FLAG-AMPKγ1. Immunoprecipitates were pre-incubated with NAD^+^ (0-6500 μM) for 1 h prior to reaction initiation. Luminescence values (RLU) were normalised to the lowest NAD^+^ concentration (0.3 μM), set to 1 for both WT and Y121A to reflect relative enzyme activity. Cells were treated with 100 nM FK for 24 h prior to lysis and immunoprecipitation. Data in L-M represent three (M) or four (L) independent experiments. **N,** Comparison of kinase activity (RLU) of immunoprecipitated WT or Y121A AMPKγ1 complexes under untreated conditions (NT), after 20 nM BAY-3827 (BAY), or with NAD^+^ at a concentration corresponding to the WT IC50 (157.8 μM) extrapolated from the curve in M. Luminescence values are shown relative to NT for each enzyme. *P* values were calculated by Ordinary one-way ANOVA and Tukey’s multiple comparisons test. \**P*<0.05; \*\**P*<0.01.

To investigate how NAD^+^ may modulate AMPK activity, we used complementary structure-based approaches including molecular docking, all-atom molecular dynamics simulations and enhanced sampling calculations. These analyses predicted that NAD^+^ can be accommodated within the γ1 CBS1 nucleotide-binding site, where its nicotinamide moiety interacts with Tyr121 (Fig. 3C, D, Fig. S3D). In this model, the adenosine moiety of NAD^+^ occupied a position similar to AMP in the reported γ1 crystal structure, with potential competition between NAD^+^ and AMP at CBS1 (Fig. 3C). Equilibrium all-atom molecular dynamics simulations showed that the Tyr121Ala (Y121A) mutation destabilised NAD^+^ binding, while having little effect on AMP association with CBS1 or on the γ1 backbone structure (Fig. 3E, F, Fig. S3F). Well-tempered metadynamics further predicted more favourable binding of NAD^+^ than AMP at CBS1, with Y121A selectively weakening NAD^+^ binding (Fig. 3G, H). Random-accelerated molecular dynamics similarly predicted a longer residence time of NAD^+^ at CBS1 than for AMP at CBS1 or CBS3, supporting prolonged NAD^+^ engagement with the predicted binding site (Fig. S3E). Together, these simulations nominated Tyr121 as a candidate determinant of NAD^+^-dependent inhibition of AMPK.

Tyr121 was conserved in vertebrates but not lower organisms, suggesting that NAD^+^ sensing by AMPK may have emerged during vertebrate evolution (Fig. 3I). Consistent with a γ-subunit-dependent interaction, Dianthus spectral shift assays *in vitro* showed stronger NAD^+^ binding to the intact AMPKα2β2γ3 holoenzyme (Kd ∼50 μM) than the isolated α2 kinase subunit or the Aurora A kinase domain (both Kd ∼ 650 μM) (Fig. S3G-I).

To experimentally test whether Tyr121 contributes to NAD^+^-dependent regulation of AMPK, FLAG-tagged wild-type and Y121A γ1 subunits were overexpressed in ARPE-19 cells and efficiently incorporated into holoenzymes with endogenous α and β subunits (Fig. S3J). Compared to wild-type γ1, the Y121A mutation rendered AMPK insensitive to NAD depletion 24 hours after FK866 treatment (Fig. 3J). At later time points, Y121A-containing AMPK was gradually activated by FK866 (Fig. 3J), likely explainable by the increase in AMP/ATP ratio observed in our metabolomics data (Fig S3K, Fig. 2A). *In vitro* kinase assays of immunoprecipitated Y121A-containing holoenzymes confirmed an approximately four-fold reduction in sensitivity to NAD^+^ inhibition compared to wild type complexes (Fig. 3K-N). Together, these data support a model in which NAD^+^ directly regulated AMPK through a γ1 Tyr121-dependent mechanism.

### Hyperactivity of AMPK due to NAD depletion suppresses autophagy

To test the role of AMPK in autophagy inhibition in response to NAD deficit, we subjected wild type and AMPKα1/α2 DKO ARPE-19 cells to FK866 treatment in the absence/presence of NR. Suppression of autophagy by FK866 was completely rescued in DKO cells based on Halo-GFP-LC3 reporter and ULK1 puncta formation (Fig. 4A, Fig. S4A, B). Next, we overexpressed wild type and Y121Aγ1 subunits in ARPE-19 cells which were subjected to the FK866/NR treatment protocol. In contrast to cells expressing wild type γ1, autophagy remained active in FK866-treated cells expressing NAD^+^-insensitive Y121A mutant (Fig. 4B). Furthermore, AMPK hyperactivation was sufficient to cause autophagy inhibition as demonstrated using AMPK activator MK-8722 whilst AMPK inhibitor BAY-3827 increased autophagy flux (Fig. 4C, Fig. S4C). Taken together, in certain cellular conditions such as NAD depletion, hyperactivation of AMPK suppresses autophagy.

**Figure 4.**
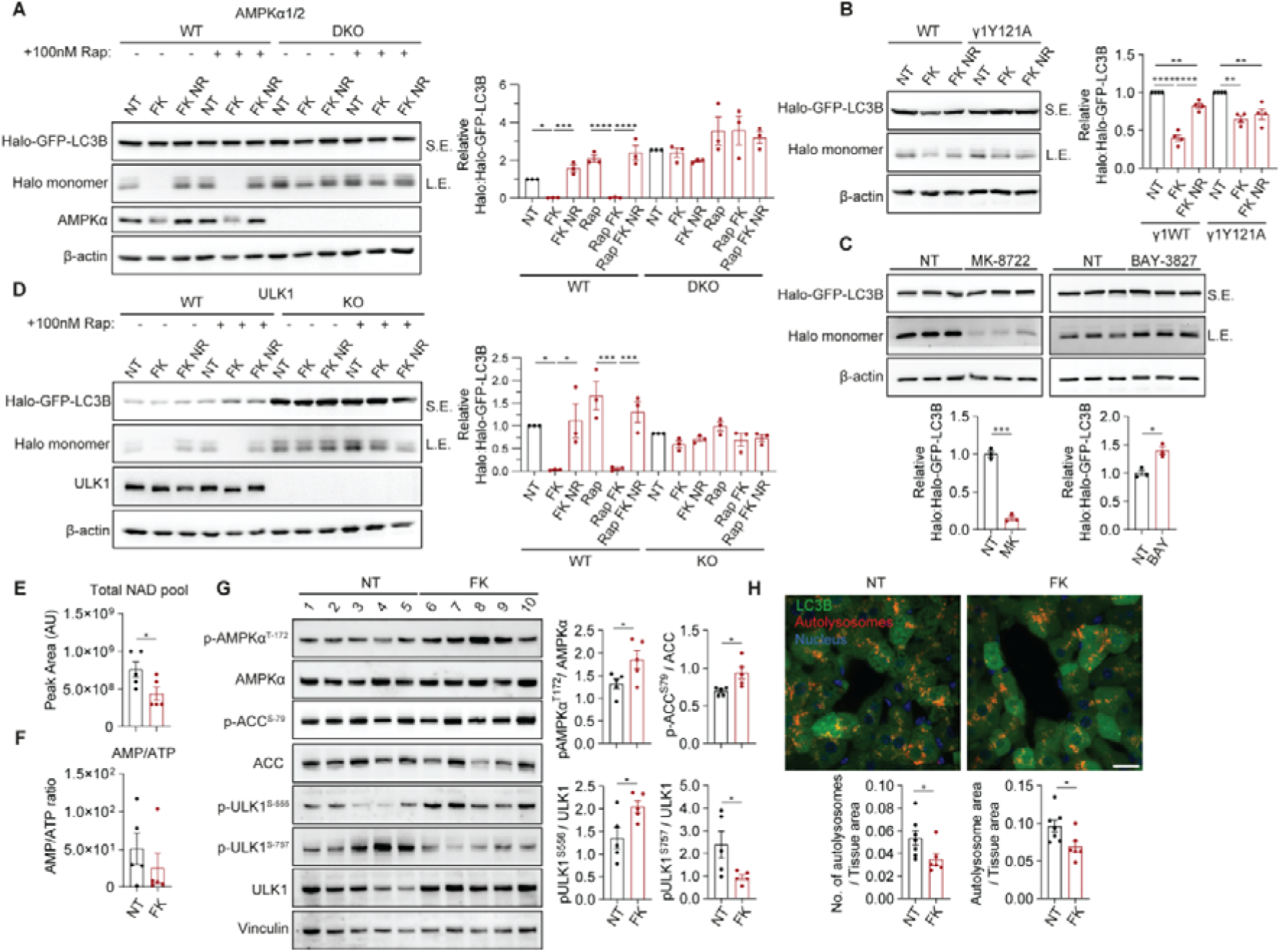
Autophagy suppression by NAD depletion requires AMPK kinase activity. **A,** Representative immunoblot and quantification of halo processing assay in AMPKα1/α2 WT or DKO ARPE-19 cells stably expressing Halo-GFP-LC3B. Cells were treated with 100 nM FK866 (FK) in the presence or absence of 1 mM nicotinamide riboside (NR), with 100 nM rapamycin (Rap) added for the final 6 h where indicated. The NT baseline of DKO was scaled to the average NT of WT to account for its higher autophagy activity. **B-D,** Representative immunoblots and quantification of the halo processing assay in ARPE-19 cells expressing Halo-GFP-LC3B, including cells stably expressing WT or Y121A FLAG-AMPKγ1 treated as in A (B), cells treated with 10 μM MK-8722 or 5 μM BAY-3827 for 72 h (C), and ULK1 WT or KO cells treated as in A (D). **E, F,** Metabolite analysis of pooled NAD^+^/NADH (E), or AMP/ATP ratio (F) in liver samples from AutoQC mice by LC-MS. Mice were dosed with vehicle or 30 mg/kg FK866 once daily for 5 days. **G,** Immunoblot analysis of indicated proteins from liver lysate of Auto-QC mice dosed as in E-F. **H,** Representative confocal micrographs and quantification of autolysosomes from liver of dosed Auto-QC mice. Data are mean ± SEM of three independent experiments (A-D), n=10 E-G (5 untreated, 5 treated) or n=13 H (7 untreated, 6 treated). *P* values were calculated by Ordinary one-way ANOVA and Tukey’s multiple comparisons test (A-B, D) or Welch’s t-test (C, E-H). \**P*<0.05; \*\**P*<0.01; \*\*\**P*<0.001; \*\*\*\**P*<0.0001. Scale bars = 20 µm (H).

Loss of ATG14 phosphorylation at an ULK1 site due to NAD depletion was also completely restored in AMPKα1/α2 DKO cells suggesting ULK1 mediates inhibitory effect of AMPK on autophagy (Fig. S4D). Indeed, defective autophagy flux was rescued in ULK1 KO ARPE-19 cells (Fig. 4D) with compensatory role of ULK2 likely to be responsible for the residual autophagy activity in ULK1 KO cells [24]. NAD depletion did not have an overt effect on the assembly of ULK1 complex suggesting phosphorylation by AMPK suppresses its activity but not structure (Fig. S4E). Collectively these data indicate that ULK1 is required for autophagy suppression downstream of AMPK during NAD depletion, with AMPK hyperactivation acting as a dominant signal that overrides mTORC1 inhibition and prevents autophagy initiation. (Fig.1,2,4).

To extend these findings to more physiologically relevant models, we first examined the effects of NAD depletion in human Embryonic Stem Cell (hESC)-derived neurons, where FK866 treatment increased AMPK activation (Fig. S4F-G). This coincided with an accumulation of p62, without overt changes to LC3B, consistent with impaired autophagy induction (Fig. S4H). To test the effect of NAD depletion on AMPK and autophagy *in vivo*, we subjected mice carrying auto-QC reporter [25] to 5 daily intraperitoneal injections of FK866 or vehicle control. The regimen was well-tolerated (at an FK866 dose of 30 mg/kg) with no significant body weight loss over the treatment duration (Fig. S4I). Upon termination of the treatment, liver tissue was subjected to metabolomics analyses, which showed a decrease in the total NAD pool (NAD^+^/NADH), without significant changes in adenylate ratios (Fig. 4E-F, Fig. S4J). Increased AMPK pathway activity and reduced ULK1 phosphorylation on the mTORC1-specific Ser757 site was identified in liver tissue from FK866-treated animals (Fig. 4G). Concomitantly, quantification of autolysosomes visualised by the auto-QC reporter indicated reduced autophagy flux following FK866 treatment (Fig. 4H).

Therefore, acute inhibition of the NAD^+^ salvage pathway both in cultured cells and *in vivo* results in the activation of AMPK which in turn suppresses the autophagy pathway. This mechanism provides potential link between the two key features of age-related disease, NAD^+^ deficit and autophagy dysfunction.

## Discussion

Here we identified AMPK as a pathway activated in conditions of NAD deficit. Functionally, this response may form part of a compensatory mechanism to restore NAD^+^ levels, as AMPK has been shown to promote NAD^+^ salvage by increasing NAMPT expression and directly phosphorylating NAMPT to enhance its activity under genotoxic stress [11,12,26]. More broadly, AMPK-dependent NAD^+^ salvage may support tissue adaptation to metabolic and environmental stresses associated with acute NAD depletion, including photodamage and inflammation [27,28]. At the same time, our data demonstrate that upon sustained NAD depletion, AMPK negatively regulates broader metabolic programmes including mTORC1 and autophagy.

Although AMPK is classically regarded as a potent activator of autophagy during energy stress, recent studies indicate a more context-dependent relationship in which AMPK can either promote or restrain autophagy dependent on the stimulus [15,18,29]. Our findings add to this complexity by showing that NAD depletion engages AMPK to suppress mTORC1 whilst also preventing autophagy activation, thereby reducing both anabolic and catabolic outputs during sustained metabolic stress. Mechanistically, this is consistent with previous work implicating the ULK1 complex as a key target through which AMPK can inhibit autophagy [15,18,30], including under conditions of mTORC1 suppression associated with amino acid starvation [15]. One possible interpretation is that under metabolic stress, AMPK-mediated restraint of energetically demanding processes, including autophagy, may conserve resources needed for immediate survival [31]. However, persistent suppression of autophagy could also compromise cellular quality control and contribute to functional decline in age-related diseases, where reduced NAD availability and impaired autophagy are both commonly implicated features. Accordingly, in addition to the previously reported role of autophagy in the maintenance of NAD [32–34], our findings suggest that NAD depletion may also contribute directly to autophagy impairment.

Regulation of AMPK by NAD-derived metabolites has previously been investigated using biochemical and biophysical approaches, but a coherent physiological model has not emerged. Early biochemical work suggested that NAD^+^ and NADH can modulate AMPK activity, whereas subsequent studies implicated reduced pyridine nucleotides, including NADH and NADPH, in competitive binding within the γ-subunit nucleotide-binding region [19–21]. However, conflicting findings and the absence of clear physiological evidence have left the functional relevance of NAD-dependent AMPK regulation unresolved [22]. Our biochemical assays support direct suppression of AMPK by NAD^+^ at physiological levels, whereas cytosolic NADH is unlikely to reach concentrations sufficient to account for the inhibitory effect observed here [35]. We therefore propose that AMPK senses NAD status primarily through NAD^+^ availability rather than through NAD redox state.

Our molecular modelling provides a potential structural basis for this mechanism, predicting NAD^+^ binding at the γ1 CBS1 site. This model is compatible with the established role of the AMPK γ-subunit as a nucleotide-binding regulatory module, and with previous biochemical and biophysical evidence that pyridine nucleotides including NADH and NADPH can bind AMPK at exchangeable γ-subunit nucleotide-binding sites [19–21]. However, no structure of AMPK bound to a pyridine nucleotide has yet been reported, leaving the structural basis of NAD-dependent regulation unresolved. Mutation of Y121, predicted to contribute to NAD^+^ binding in our model, reduced the sensitivity of AMPK to NAD depletion, identifying this region as a candidate determinant of NAD-dependent regulation. Together, these findings provide a starting point for future biochemical and structural studies of this regulatory switch. Finally, the evolution of NAD sensing by AMPK represents an intriguing direction for future research, particularly if this potentially vertebrate-specific mechanism contributes to tighter metabolic control, stress adaptation and longevity.

### Limitations of the study

Although our findings provide evidence that AMPK can sense NAD availability in addition to classical AMP/ADP/ATP ratios, this report opens numerous questions for future studies. At the molecular level, it remains unclear how NAD^+^ binding at the γ1 CBS1 site inhibits AMPK beyond the predicted interference with AMP-dependent allosteric activation. Experimental structural evidence will be required to validate this model and define the molecular basis of NAD-dependent regulation. At organismal level, previous study employing *Nampt* KO mouse model did not report overt changes in AMPK or other metabolic pathways, potentially indicating existence of biological safety checks ensuring adaptation to chronic NAD deficit [36]. Therefore, the physiological relevance and temporal delineation of NAD depletion in cellular and tissue metabolic homeostasis, as well as in ageing and age-related disease requires further exploration. Such studies may help identify whether targeting NAD-dependent AMPK signalling can improve stress resistance and limit age-related functional decline.

## Acknowledgements

We are grateful to Newcastle Bioimaging Unit for imaging assistance and Richard Heath for advice. This work is supported by grants from Medical Research Council (MR/Z504488/1) to S.S. and V.I.K.; (MC_UU_00038/2) to I.G.G.; Action Medical Research/LifeArc (GN3049), LifeArc (Philanthropic Fund P2019-0004, and Pathfinder Award), Wolfram Syndrome UK, and Birmingham Fellowship to S.S.; Funding from the Novo Nordisk Foundation (NNF23SA0084103) to K.S.; JSPS (20K22912; 25K18725), AMED (JP24gm6710024), LOTTE Foundation, Nakajima Foundation, Uehara Memorial Foundation, Senri Life Science Foundation and Sumitomo Foundation, and Nippon Shinyaku to T.K. (the funder Nippon Shinyaku was not involved in the study design, collection, analysis, interpretation of data, the writing of this article, or the decision to submit it for publication); a Wellcome Trust Senior Fellowship 222531/Z/21/Z to E.Z.; grants from a BBSRC AGENT Network (BB/W018381/1), VitaDAO/Molecule and Procter & Gamble academic partnerships to V.I.K.; and a China Scholarship Council-Newcastle University studentship to J.T. and V.I.K. Both S.S. and V.I.K. are Former Fellows for life at Hughes Hall, University of Cambridge, United Kingdom.

**Niall Wilson:** Formal analysis; investigation; visualisation; methodology; writing – original draft; writing – review and editing. **Yoana Rabanal Ruiz:** Formal analysis; investigation; visualisation; methodology; writing – review and editing. **Aniketh Bishnu:** Formal analysis; investigation; visualisation; methodology. **Shangze Xu:** Formal analysis; investigation; visualisation; methodology. **Congxin Sun:** Formal analysis; investigation; visualisation; methodology. **Mohd Sayed Ahangar:** Formal analysis; investigation; visualisation; methodology. **Kevin M. Rattigan:** Formal analysis; investigation; methodology. **Jiangyu Tang:** Formal analysis; investigation. **Danyelle Silva-Amaral:** Formal analysis; investigation. **Conchita Fraguas Bringas:** Formal analysis; investigation **Tetsushi Kataura:** Formal analysis; investigation; writing – review and editing. **Sovan Sarkar:** Supervision; resources; writing – review and editing. **Kei Sakamoto:** Supervision; resources. **Elton Zeqiraj:** Supervision; resources; writing – review and editing. **G. Vignir Helgason:** Supervision; resources; writing – review and editing. **Ian G. Ganley:** Supervision; resources; writing – review and editing. **Agnieszka K. Bronowska:** Supervision; investigation; visualisation; writing – review and editing. **Viktor I Korolchuk:** Conceptualisation; supervision; visualisation; funding acquisition; project administration; writing – original draft; writing – review and editing.

## Inclusion and Diversity

We support inclusive, diverse, and equitable conduct of research.

## STAR Methods

### RESOURCE AVAILABILITY

#### Lead contact

Further information and requests for resources and reagents should be directed to and will be fulfilled by the lead contact, Viktor I. Korolchuk.

#### Materials availability

Further information and requests for resources and reagents listed in Key Resources Table should be directed to the Lead Contact.

#### Data and code availability

- Uncropped immunoblot images have been deposited at Mendeley and are publicly available as of the date of publication. The DOI is listed in the key resource table. Other data reported in this paper will be shared by the lead contact upon request.
- This paper does not report original code.
- Any additional information required to reanalyse the data reported in this paper is available from the lead contact upon request.

### EXPERIMENTAL MODEL AND STUDY PARTICIPANT DETAILS

#### Cell culture

Human ARPE-19 retinal pigment epithelia cells were cultured in DMEM: F12 media (Gibco) supplemented with 10% foetal bovine serum, 100 U/ml penicillin/streptomycin in atmospheric oxygen conditions at 37LC and 5% CO_2_ in a humidified incubator. ARPE-19 AMPKα1/α2 WT / DKO and FLAG-ULK1 KO / WT were generated in the previous study [18]. HMC3 cells were maintained in high-glucose DMEM (Gibco), supplemented with 10% FBS, 100 U/ml penicillin/streptomycin and 2 mM L-glutamine at 37LC and 5% CO_2_.

#### Culture of human ESC

WIBR3 human embryonic stem cells (hESCs) were maintained under feeder-free conditions on Geltrex basement membrane matrix-coated plates in StemFlex Basal Medium supplemented with StemFlex 10X Supplement and 1 % penicillin/streptomycin (all from Gibco), at 37 °C in a humidified incubator containing 5 % CO2 and 5 % O2 [34,37]. The hESCs were differentiated into neural precursors (NPs), as described previously [34]. The NPs were cultured on Poly-L-ornithine and Laminin (PO-L) (Sigma-Aldrich) coated plates or flasks in medium comprised of DMEM/F-12 medium containing 1 % N-2 supplement, 2 % B-27 supplement, 1% penicillin/streptomycin, 1 % non-essential amino acids, 1 % L-Glutamine (all from Gibco), supplemented with 10 ng/mL FGF-2 (Miltenyi Biotec) and 10 ng/mL EGF (PeproTech), and maintained in a humidified incubator with 5 % CO2 at 37 °C. The NPs were further differentiated into neurons for 4 weeks, as described previously [34]. Neuronal differentiation was done using medium comprised of DMEM/F-12 medium containing 1 % N-2 supplement, 2 % B-27 supplement, 1% penicillin/streptomycin, 1 % non-essential amino acids, 1 % L-Glutamine (all from Gibco), and maintained in a humidified incubator with 5 % CO2 at 37 °C. For NAD depletion experiments, hESC-derived neurons were treated with DMSO (vehicle control) or 10 nM FK866 (NAMPT inhibitor) for the last 6 days of the 4-weeks neuronal differentiation period.

#### Animal model

Auto-QC mice expressing the mCherry-GFP-LC3B reporter were generated as previously described [25]. Approximately 3-month-old mice of both sexes were used for all experiments and were housed in sex-match groups of two to five animals under standard conditions (12 h light/dark cycle, 21±10°C, 55-65% relative humidity). Food and water were provided *ad libitum*.

Mice were randomly assigned to experimental groups and administered FK866 via intraperitoneal (IP) injection at a dose of 30 mg/kg, dissolved in DMSO and diluted in 0.9% saline, or vehicle control, once daily for five consecutive days (vehicle, n=7; FK866, n=6). Animals were euthanised approximately 4 h after the final dose. Liver tissue was collected and immediately processed for microscopy and immunoblot analysis. All animal experiments were approved by the University of Dundee ethical review committee and conducted in accordance with UK Home Office regulations and the Animals Act 1986.

## METHOD DETAILS

### Plasmids

Plasmids encoding 3x FLAG-tagged AMPK γ1WT and γ1Y121A were generated by VectorBuilder using the PRKAG1 transcript variant 1 mRNA (NM_002733.5) sequence. pMRX-No-HaloTag7-mGFP-LC3B was a gift from Noboru Mizushima, Addgene, 184901) [38]. Further information on plasmids used in this study can be found in Table 5.

### Viral production and generation of stable cell lines

Generation of ARPE-19 and HMC3 cells stably expressing AMPK γ1WT or γ1Y121A, and Halo-GFP-LC3B was achieved by packaging lentiviruses in the HEK293FT (293FT) cell line. 293FT cells were seeded into a 10 cm dish (6.0×10^6^ cells/10 mL/dish) in antibiotic-free glucose culture medium. Next day, cells were transfected with plasmids containing the packaging psPAX2 (gift from Didier Trono) and envelope pCMV-VSV-G genes (gift from Bob Weinberg) [39], and the pCMV-FLAG-AMPK γ1WT/Y121A and pMRS-No-HaloTag7-LC3B constructs with Lipofectamine™ 3000 Transfection Reagent (Invitrogen). Following overnight transfection, the medium was replaced with fresh antibiotic-free medium that was collected after 24 h. Virus-containing medium was filtered through a 0.45 µm pore-size filter and overlaid on 30% confluent cells for 24 h in the presence of 10 µg/ml polybrene. Cells stably expressing the AMPK γ1WT or γ1Y121A subunit were optimised for protein expression via 2 μg/mL puromycin selection for 7 days, and equal expression between lines was monitored by anti-FLAG immunoblotting.

### Immunoblot analysis

Immunoblotting was performed as previously described [40]. In brief, cell lysates were collected in RIPA buffer (Sigma-Aldrich) supplemented with 2x Halt™ protease and phosphatase inhibitor cocktail (Thermo Fisher Scientific). Protein concentration was determined using the DC Protein Assay (Bio-Rad) and samples were prepared by boiling for 5 min at 100LC in Laemmli sample buffer (Bio-Rad) in the presence of 2.5% β-mercaptoethanol. Equal amounts of protein (20 – 40 μg) were subjected to SDS-PAGE and transferred to PVDF membranes. Membranes were blocked in 5% milk (Merck Millipore) or 5% BSA (Sigma-Aldrich) in PBS with 0.1% [v/v] Tween® 20 (Sigma-Aldrich) for 1 h at room temperature and incubated with primary antibodies overnight at 4LC. Secondary antibodies conjugated to horseradish peroxidase (HRP) for rabbit and mouse were all used at 1:5000 dilution for 1 h at room temperature. Chemiluminescence detection was achieved using Clarity Western ECL Substrate (Bio-Rad) and an iBright™ CL1500 Imaging System (Invitrogen). Densitometry analyses of immunoblots were conducted using Fiji/ImageJ (version 1.48, National Institutes of Health). Unless otherwise stated, quantified data was normalised to the untreated (NT) group. See the Table 1 for information of antibodies used for immunoblotting.

**Table 1.** Immunoblot primary and secondary antibodies.

| Target | Species | MW (kDa) | Dilution | Vendor | Cat. No. |
| --- | --- | --- | --- | --- | --- |
| β-Actin | Mouse | 49 | 1:4000 | St. John's Laboratory | |
| Vinculin | Mouse | 124 | 1:4000 | Sigma-Aldrich | V9131 |
| p-AMPKα (Thr172) | Rabbit | 62 | 1:1000 | CST | 2535 |
| AMPKα | Rabbit | 62 | 1:1000 | CST | 5831 |
| p-ACC (Ser79) | Rabbit | 280 | 1:1000 | CST | 11818 |
| ACC | Rabbit | 280 | 1:1000 | CST | 3676 |
| p-ULK1 (Ser555/6) | Rabbit | 150 | 1:1000 | CST | 5869 |
| p-ULK1 (Ser757) | Rabbit | 150 | 1:1000 | CST | 6888 |
| ULK1 | Rabbit | 150 | 1:1000 | CST | 8054 |
| p-S6 (Ser235/236) | Rabbit | 32 | 1:1000 | CST | 4858 |
| S6 | Rabbit | 32 | 1:1000 | CST | 2217 |
| NAMPT | Rabbit | 52 | 1:1000 | CST | 86634 |
| p-S6K (Thr389) | Rabbit | 70, 85 | 1:1000 | CST | 9234 |
| S6K | Rabbit | 70, 85 | 1:1000 | CST | 2708 |
| p-TSC2 (Ser1387) | Rabbit | 200 | 1:1000 | CST | 5584 |
| TSC2 | Rabbit | 200 | 1:1000 | CST | 4308 |
| p-AKT (Thr308) | Rabbit | 60 | 1:1000 | CST | 13038 |
| AKT | Rabbit | 60 | 1:1000 | CST | 9272 |
| p-mTOR (Ser2448) | Rabbit | 289 | 1:1000 | CST | 5536 |
| mTOR | Rabbit | 289 | 1:1000 | CST | 2983 |
| LC3B | Rabbit | 14, 16 | 1:1000 | CST | 3868 |
| LC3B | Rabbit | 14, 16 | 1:2000 | Novus Biologicals | NB100-2220 |
| HaloTag | Rabbit | 34 (monomer) | 1:1000 | Promega | G9211 |
| ATG13 | Rabbit | 72 | 1:1000 | CST | 13468 |
| FIP200 | Rabbit | 200 | 1:1000 | CST | 12436 |
| p-ATG14 (Ser29) | Rabbit | 65 | 1:1000 | CST | 92340 |
| ATG14 | Rabbit | 65 | 1:1000 | CST | 96752 |
| p-ATG16L1 (Ser278) | Rabbit | 68 | 1:1000 | CST | 45511 |
| ATG16L1 | Rabbit | 66, 68 | 1:1000 | CST | 8089 |
| p-Beclin-1 (Ser30) | Rabbit | 60 | 1:1000 | CST | 54101 |
| Beclin-1 | Rabbit | 60 | 1:1000 | CST | 3738 |
| Acetyl-lysine (Ack) | Rabbit | Pan | 1:1000 | CST | 9441 |
| Poly/mono-ADP-Ribose (PAR) | Mouse | Pan | 1:1000 | Enzo Life Sciences | ALX-804-220-R100 |
| AMPKβ1/2 | Rabbit | 30, 38 | 1:1000 | CST | 4150 |
| P62 | Mouse | 62 | 1:1000 | Waters Biosciences | 610832 |
| FLAG (DYKDDDDK) | Mouse | N/A | 1:1000 | CST | 14793 |
| Anti-Mouse IgG– Peroxidase antibody | Goat | N/A | 1:5000 | Sigma-Aldrich | A2554 |
| Anti-Rabbit IgG– Peroxidase antibody | Goat | N/A | 1:5000 | Sigma-Aldrich | A0545 |
| Goat Anti-Mouse IgG, H&L Chain Specific Peroxidase Conjugate | Goat | N/A | 1:5000 | Calbiochem | 401253 |
| Goat Anti-Rabbit IgG, H&L Chain Specific Peroxidase Conjugate | Goat | N/A | 1:5000 | Calbiochem | 401393 |

### Immunoprecipitation

All immunoprecipitations in this study used Anti-FLAG® M2 Magnetic Beads (Merck Millipore). Cells were lysed in lysis buffer (50 mM Tris-HCl, pH 7.4; 150 mM NaCl; 1 mM EDTA; 1% Triton™ X-100), supplemented with 2x Halt™ protease and phosphatase inhibitor cocktail. Cell lysates were incubated with 15 µl (per 10-cm dish) magnetic FLAG antibody-coated beads overnight at 4°C with constant rotation. Following incubation, beads were precipitated in a magnetic rack and washed three times in TBS wash buffer (50 mM Tris HCl, pH 7.4; 150 mM NaCl). Proteins were eluted using 30 µl 2X Laemmli sample buffer, by heating to 100°C for 5 min. Eluates were centrifuged at 16,100L*g* for 3 min at room temperature to separate the beads before being subjected to Western blot analysis.

### Immunofluorescence

Immunofluorescence analyses were performed as described previously [40]. In brief, cells were seeded onto coverslips in 24-well plates and fixed in 4% paraformaldehyde in PBS for 10 min at room temperature followed by permeabilisation in 0.5% Triton X-100 (Sigma-Aldrich) in PBS for 10 min at room temperature. Cells were then blocked for 1 h in 5% normal goat serum (Sigma-Aldrich) in PBS with 0.05% [v/v] Tween® 20 at room temperature and incubated with primary antibodies overnight at 4°C. Primary antibodies were used at 1:200. Cells were washed three times in PBS and incubated with the appropriate secondary antibodies conjugated with Alexa Fluor 594 for 1 h at room temperature. Cells were washed, and coverslips were mounted on slides with ProLong™ Gold antifade mountant with DAPI (Invitrogen). Fluorescence images were obtained using a Leica DMi8 inverted microscope with a Plan-Apochromat 63x/1.30 oil immersion objective, equipped with an ORCA-Flash4v2.0 camera (Hamamatsu). Images were analysed in Fiji/ImageJ (version 1.48; National Institutes of Health). See Table 2 for further information on antibodies used for immunofluorescence in this study.

**Table 2.** Immunofluorescence primary and secondary antibodies.

| Target | Species | Dilution | Vendor | Cat. No. |
| --- | --- | --- | --- | --- |
| ULK1 | Rabbit | 1:100 | CST | 8054 |
| ATG13 | Rabbit | 1:100 | CST | 13468 |
| FIP200 | Rabbit | 1:100 | CST | 12436 |
| ATG16L1 | Rabbit | 1:100 | Abcam | ab187671 |
| TUJ1 (TUBB3) | Mouse | 1:200 | BioLegend | 801201 |
| p-AMPKα Thr172 | Rabbit | 1:100 | CST | 2535 |
| Goat anti-rabbit IgG(H+L), AF594 | Goat | 1:500 | Invitrogen | A-11012 |
| Goat anti-mouse IgG(H+L), AF594 | Goat | 1:500 | Invitrogen | A-11005 |
| Goat anti-rabbit IgG(H+L), AF488 | Goat | 1:500 | Invitrogen | A-11008 |

**Table 3.** Reagents, consumables and chemicals.

| Reagent | Vendor | Catalogue No. |
| --- | --- | --- |
| RIPA buffer | Sigma-Aldrich | R0278 |
| Halt™ protease and phosphatase inhibitor cocktail (100x) | Fisher Scientific | 1861280 |
| DC Protein Assay Kit | Bio-Rad | 500-0112 |
| 2x Laemmli Sample Buffer | Bio-Rad | 1610737 |
| 4x Laemmli Sample Buffer | Bio-Rad | 1610747 |
| β-mercaptoethanol | Sigma-Aldrich | M3148 |
| Empty Gel Cassettes, mini, 1.0mm | Invitrogen | NC2010 |
| Empty Gel Cassettes, mini, 1.5mm | Invitrogen | NC2015 |
| Novex Tris-glycine 4-20% gradient gel, mini, 15-well | Invitrogen | 15507460 |
| 30% Acrylamide/Bis-acrylamide | Thistle Scientific | 20-2100-10 |
| Ammonium persulfate | Sigma-Aldrich | A3678 |
| Trizma® base (Tris) | Sigma-Aldrich | T1503 |
| Glycine | Sigma-Aldrich | G8898 |
| SDS 20% |  | 10607443 |
| Sodium Chloride | Sigma-Aldrich | S7653 |
| N,N,N',N'-Tetramethylethylenediamine (TEMED) | Sigma-Aldrich | T9281 |
| Isopropanol | Sigma-Aldrich | 278475 |
| Precision Plus Protein™ Dual Colour Protein Standards |  | 610374 |
| Polyoxyethylene 20 (Tween® 20) | Sigma-Aldrich | 93774 |
| Methanol | Sigma-Aldrich | 32213 |
| Thick blotting pads | VWR | 732-0594 |
| Immobilon-P polyvinylidene difluoride (PVDF) membrane | Millipore | IPVH00010 |
| Skim Milk Powder | Sigma-Aldrich | 70166 |
| Bovine Serum Albumin (BSA) | Sigma-Aldrich | A2153 |
| Clarity Western ECL Substrate |  | 170-5061 |
| 20X PBS | CST | 9808 |
| Trichloroacetic acid (TCA) | Sigma-Aldrich | T4885 |
| Sodium hydroxide (NaOH) | Sigma-Aldrich | S5881 |
| EDTA | Fisher Scientific | BP120-500 |
| Sodium phosphate, Dibasic (Na<sub>2</sub>HPO<sub>4</sub>) | Sigma-Aldrich | 567550 |
| Tris hydrochloride (Tris-HCl) | Sigma-Aldrich | 10812846001 |
| Resazurin sodium salt | Sigma-Aldrich | R7017 |
| Riboflavin 5'-monophosphate sodium salt (FMN) | Sigma-Aldrich | F8399 |
| Ethanol | Sigma-Aldrich | 493511 |
| Alcohol Dehydrogenase | Sigma-Aldrich | A7011 |
| Diaphorase | Sigma-Aldrich | D5540 |
| β-NAD<sup>+</sup> | Sigma-Aldrich | 10127695001 |
| 10cm TC plates (TPP) | Helena Biosciences | 93100 |
| 0.45 μm Membrane Filter, PTFE (Sterile) | Starlab | P7166-6800 |
| Polybrene | Sigma-Aldrich | H9268 |
| Puromycin dihydrochloride | Fisher Scientific | A1113803 |
| MEM Non-essential Amino Acid Solution (100x) | Gibco | 11140-035 |
| OptiMEM | Invitrogen | 11058021 |
| Lipofectamine® 2000 transfection reagent | Invitrogen | 10696153 |
| Human EGF recombinant protein | Gibco | PHG0313 |
| Human FGF-basic recombinant protein | Invitrogen | RP-8627 |
| B-27 Supplement | Gibco | 12587-010 |
| Geltrex | Gibco | A1413302 |
| Mouse laminin | Sigma-Aldrich | L2020 |
| N-2 supplement (100X) | Gibco | 17502-048 |
| Poly-L-Ornithine | Sigma-Aldrich | P4957 |
| ProLong Gold Antifade Mountant with DAPI | Invitrogen | P36931 |
| StemFlex Basal Medium | Gibco | A3349401 |
| StemFlex 10X Supplement | Gibco | A3349201 |

**Table 4.**
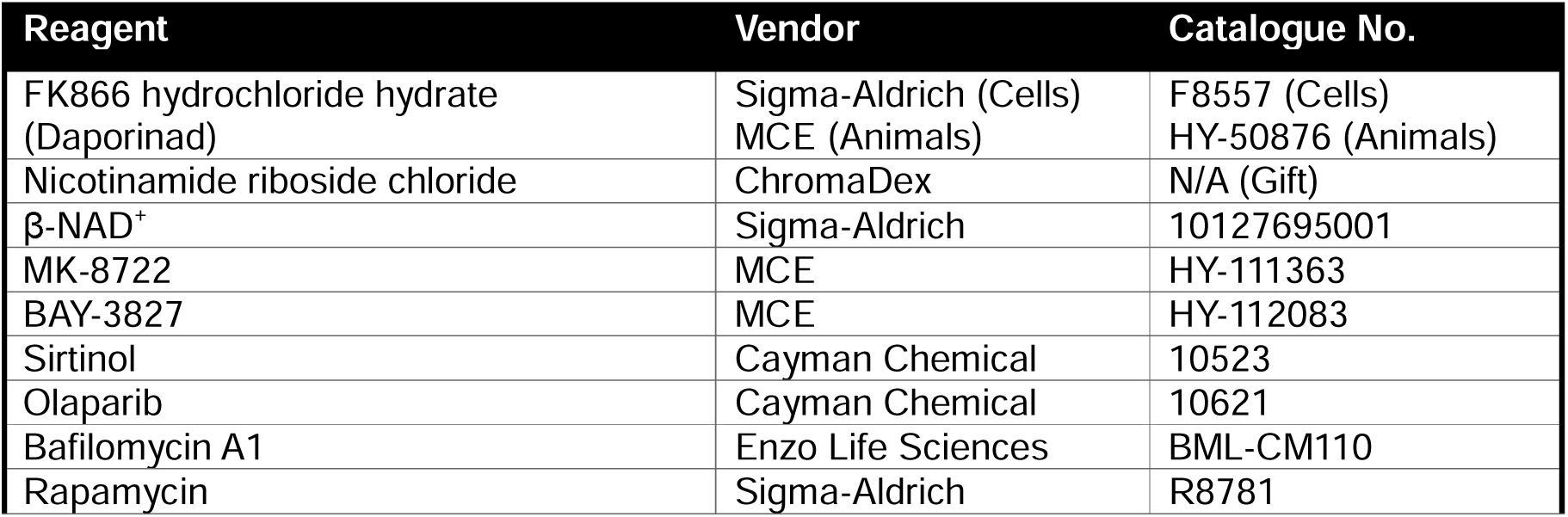
Treatments.

| Reagent | Vendor | Catalogue No. |
| --- | --- | --- |
| FK866 hydrochloride hydrate (Daporinad) | Sigma-Aldrich (Cells)<br>MCE (Animals) | F8557 (Cells)<br>HY-50876 (Animals) |
| Nicotinamide riboside chloride | ChromaDex | N/A (Gift) |
| β-NAD<sup>+</sup> | Sigma-Aldrich | 10127695001 |
| MK-8722 | MCE | HY-111363 |
| BAY-3827 | MCE | HY-112083 |
| Sirtinol | Cayman Chemical | 10523 |
| Olaparib | Cayman Chemical | 10621 |
| Bafilomycin A1 | Enzo Life Sciences | BML-CM110 |
| Rapamycin | Sigma-Aldrich | R8781 |

**Table 5.** Plasmids.

| Plasmid | Source | Catalogue No. | Gifted by |
| --- | --- | --- | --- |
| pMRX-No-HaloTag-mGFP-LC3B | Addgene | 184901 | Noboru Mizushima, University of Tokyo, Japan |
| pLV-CMV-FLAG-AMPKγ1WT-IRES-Puro | VectorBuilder | VB241103-1050paq | This study |
| pLV-CMV-FLAG-AMPKγ1Y121A-IRES-Puro | VectorBuilder | VB241103-1051rgh | This study |

### Autophagy assays

LC3B turnover assay was performed to monitor autophagy flux in ARPE-19 and HMC3 cells[41]. Cells were seeded into 12 well plates and treated with appropriate compounds for 72 h. For the final 6 h of treatment, cells were treated with bafilomycin A1 (BafA1) (to block lysosomal degradation) and were subjected to immunoblotting. LC3B-II flux was expressed by subtracting the signals of LC3B in BafA1-untreated conditions from those of LC3B in BafA1-treated conditions.

For western blot analysis of autophagy activity by halo processing assay, cells stably expressing Halo-GFP-LC3B were seeded into 6 well plates and treated with appropriate compounds for 72 h. For the final 6 h of treatment, cells were treated with 20 µM 7-Bromo-1-heptanol without media change. Lysates were subjected to immunoblot analysis using Halo antibody. Autophagic activity was quantified as a ratio of Halo monomer per Halo-GFP-LC3B.

To assess autophagy by imaging, cells expressing Halo-GFP-LC3B were seeded onto coverslips in 24-well plates and treated as above. For the final 6 h of treatment, cells were treated with 40 nM Janelia Fluor 646 Halo ligand. Cells were washed twice in PBS, fixed in 4% paraformaldehyde in PBS, and mounted onto slides as described above. Fluorescence images were obtained using an inverted Leica DMi8 microscope with a Plan-Apochromat 63x/1.40 oil immersion lens, equipped with an ORCA-Flash4v2.0 camera (Hamamatsu). Images were analysed in Fiji/ImageJ (version 1.48; National Institutes of Health), and quantification was performed on at least 50 cells per condition. The number of autophagosomes (GFP^+^ JF646^+^ puncta) and autolysosomes (GFP^-^ JF646^+^ puncta) per cell were quantified by outlining single cells as regions of interest.

### *In vitro* AMPK kinase assay

AMPK activity was measured *in vitro* using the AMPK (A1/B1/G1) Kinase Enzyme System coupled with the ADP-Glo^TM^ Kinase Assay (Promega, V9021), in white 384-well microplates (Fisher Scientific, 164610). Recombinant full-length His-tagged human AMPKα1β1γ1 (Sino Biological, P47-10H) was diluted in kinase buffer (40 mM Tris, pH 7.5; 20 mM MgCl_2_; 0.1 mg/ml BSA; 50 µM DTT) to a final amount of 10 ng per well, determined by initial enzyme titration experiments.

The enzyme was incubated for 1 h at room temperature in the presence or absence of BAY-3827, NAD^+^ (diluted in H_2_O), and/or 100 µM AMP. Tool compound concentrations are indicated in figure legends, and the final DMSO concentration for all assays was maintained at 2%. Following compound incubation, SAMStide peptide substrate (HMRSAMSGLHLVKRR, 1 µg/µL) and ATP (50 µM) were added to initiate the kinase reaction, resulting in a final reaction volume of 7.25 µL. Reactions were incubated for 1 h at room temperature. To terminate the kinase reaction and convert remaining ATP to ADP, an equal volume (7.25 µL) of ADP-Glo^TM^ reagent was added to each well and incubated for 40 min at room temperature. Next, 14.9 µL Kinase Detection Reagent was added and incubated for a further 30 min to generate the luminescent signal. Luminescence, corresponding to ADP production during the kinase reaction, was measured using a FLUOstar Omega microplate reader (BGM Labtech). All assay steps were performed under minimal light exposure. Where applicable, luminescence values were normalised by subtracting the background signals from no-enzyme control wells.

For detecting kinase activity of AMPK complexes containing FLAG-γ1 WT or Y121A, cell lysates were subjected to anti-FLAG immunoprecipitation as described above. Both WT and Y121A samples were prepared under equivalent conditions. Following overnight incubation of 1 mg of total protein at 4°C with Anti-FLAG® M2 Magnetic Beads, the unbound fraction was discarded, and the beads were washed in TBS wash buffer (50 mM Tris HCl, pH 7.4; 150 mM NaCl). Beads were then resuspended in storage buffer (10 mM sodium phosphate, pH 7.4, adjusted with HCl, 150 mM NaCl, 50% [v/v] glycerol) and stored at-80°C until use. For kinase activity assays, bead suspensions were then diluted in dH_2_O (16.6% [v/v]) and loaded directly into wells. Assays were performed using the same protocol described for the recombinant AMPK kinase assay. Luminescence values were normalised to empty bead controls (no enzyme bound).

### NAD^+^ and NADH measurements

Measurements of NAD^+^ and NADH in mammalian whole-cell lysates were performed as described in a previously published protocol [32,42]. NAD^+^ or NADH was extracted from 4× 10^6^ cells (NAD^+^) or 8×L10^6^ cells (NADH) cells by probe sonication with an acidic solution (10% trichloroacetic acid (TCA) (Sigma-Aldrich) or basic solution (0.5 M sodium hydroxide (Sigma-Aldrich), 5L□M EDTA (Sigma-Aldrich)) respectively. NADH samples were heated at 60 °C for 30□min. Samples were centrifuged at 16,100L*g* for 3□min at 4 °C. Supernatants were collected and 10% volume of 1□M Tris (Sigma-Aldrich) was added to adjust pH, followed by NAD^+^ and NADH measurements. Aliquots of supernatant (NADH) or the pellet (NAD^+^) resolved in 0.2 M sodium hydroxide were used to measure protein concentrations. NAD^+^ and NADH levels were determined by the fluorescence intensity of resorufin produced by an enzymatic cycling reaction using resazurin, riboflavin 5′-monophosphate, alcohol dehydrogenase, and diaphorase (all from Sigma-Aldrich). Fluorescence intensity was monitored every minute for a total 60□min using a microplate reader (FLUOstar Omega, BMG Labtech). NAD^+^ and NADH levels were determined by a β-NAD (Sigma-Aldrich) standard curve and adjusted to protein concentration determined by the DC protein assay (BioRad).

### LC-MS-based metabolomics

Metabolites were isolated from ARPE-19 cells following an established protocol described previously [33]. Cells were maintained in 10 cm dishes under routine culture conditions and treated as specified in the figure legends. Samples were collected at ∼85% confluency. Cells were rinsed twice in ice-cold PBS and lysed at a density of 2×106 cells/mL in a metabolite extraction buffer composed of 50% methanol (Fisher Scientific), 30% acetonitrile (Sigma-Aldrich), and 20% deionised water. Samples were vortexed for 45 s and centrifuged at 16,100 *g* for 10 min at 4°C. Supernatants were collected and analysed by LC-MS using a three-point calibration curve with uniformly labelled 13C/15N amino acids for quantification. Extraction buffer blanks were processed and analysed in parallel.

Samples were analysed by LC-MS using an UltiMate 3000 HPLC system (Fisher Scientific) coupled to a qExactive Plus Orbitrap mass spectrometer (Thermo Scientific). Metabolite separation was performed on a ZIC-pHILIC column (150 x 2.1 mm, 5 µm; Merck) with a matching guard column, using 20 mM ammonium carbonate containing 0.1% ammonium hydroxide in water (A) and acetonitrile (B) as mobile phases. A linear biphasic gradient from 80% to 20% (mobile phase B) over 15 min was applied at a flow rate of 200 µL/min, followed by equilibration, giving a total run time of 22 min, with the column maintained at 45°C. MS data were acquired in polarity-switching mode following calibration with a custom CALMIX solution. Full-scan MS1 spectra were collected in profile mode over an m/z range of 75-1000 at a resolution of 70,000, with an AGC target of 1×106 and a maximum injection time of 250 ms. Source parameters were set as specified [43], and data were acquired using Xcalibur software, achieving mass accuracy below 5 ppm. Peak detection and quantification were performed by the collaborators using Skyline (v24.1, UoW, Seattle, USA). Metabolite identities were assigned using an in-house reference library compiled from KEGG, HMDB and LIPID MAPS, based on accurate mass matching and known retention times on the pHILIC column. Accurate ion masses and retention times has been previously established on the LC-MS system using a commercially available metabolite standard mix (Merck: MSMLS-1EA) [43]. Data was subsequently processed into charts using GraphPad Prism.

Mouse tissue metabolomics was performed under the same extraction and analytical conditions described above using pulverised cryo-frozen liver samples from AutoQC mice included in this study. For each sample, 20 mg tissue was extracted in 1000 µL extraction buffer, in the absence of stable isotope labelling using stainless steel beads and a TissueLsyer II (Qiagen) (2 x 2 min pulses at 30 Hz with 2 min storage at-20 °C between pulses. Data were corrected by loading of a blank solvent buffer control sample. Due to co-elution and potential for *ex-vivo* interconversion of NAD^+^ and NADH, the total NAD pool (NAD^+^ and NADH together) was used for primary interpretation of this metabolite group, although individual NAD^+^ and NADH measurements were also analysed separately as supplement.

### siRNA transfection

Knockdown of human *NAMPT* was performed by siRNA transfection. ON-TARGETplus SMARTpool siRNA against human *NAMPT* (L-004581-00-0010) *was* purchased from Horizon Discovery. Final siRNA concentrations of 20 or 100 nM were used for silencing, and transfections were performed using Lipofectamine 2000 (Invitrogen) as per manufacturer’s instructions. Briefly, ARPE-19 cells were seeded into a 6-well plate and cultured for 24 h, and then transfected with 100 nM of siRNA with Lipofectamine 2000 reagent for 24 h. Cells were then passaged and cultured for 24 h, followed by a second transfection with 20 nM siRNA and 72 h treatment with NR. See Table 6 for further information on the siRNAs used in this study.

**Table 6.** siRNA.

| siRNA | Accession hits | Catalogue No. |
| --- | --- | --- |
| SMARTpool: ON-TARGETplus human NAMPT siRNA | NM_005746 | L-004581-00-0010 |
|  | XM_047419699 |  |
|  | XM_047419700 |  |

### Molecular docking

Molecular docking was performed in SeeSAR (v. 14.2) [44], using the 2.5 Å human AMPK heterotrimer bound to the allosteric activator C455 (PDB: 8BIK) [45]. Putative binding sites were identified with the SeeSAR Site module, followed by non-covalent docking of NAD^+^, AMP and control allosteric inhibitors using up to 500 poses per run, ring puckering, high clash tolerance and fixed ligand chirality. For the AMPK γ1 subunit, docking was repeated in template mode using AMP as the template. Binding affinities were estimated with the HYDE [46] scoring function, which incorporates intrinsic protein-ligand interactions and desolvation terms.

### All-atom molecular dynamics (MD) simulations

Complexes for MD simulations were prepared in UCSF Chimera [47]. Water molecules, non-complexes ions and crystallisation additives were removed; missing hydrogens and incomplete side chains were asses using the Dunbrack rotamer library [48], and alternative side-chain conformations were resolved by retaining the highest occupancy conformer. The Y121A mutation was generated in UCSF Chimera. NAD^+^ and AMP were parameterised using ACPYPE [49] with the GAFF force field [50] and AM1-BCC partial charges [51]. Additional force-field definitions were generated using an in-house module implemented in the GMX ToolBox webserver.

MD simulations were performed in GROMACS 2023.2 [52]. The AMPK γ1 subunit was parameterised with the AMBER99SB-ILDN force field and solvated with the TIP3P water model [53]. Systems were placed in a periodic box with a 1 nm buffer, solvated, neutralised and supplemented with NaCl to 0.1 M. After energy minimisation, systems were heated to 300 K for 20 ps under backbone restrains in the NVT ensemble, followed by 1 ns NPT equilibration at 300 K and 1 bar. Long-range electrostatics were treated using Particle Mesh Ewald [54], bond were constrained using LINCS, and pressure was maintained with the Parrinello-Rahman barostat [55]. Production simulations were run for 500 ns in the NPT ensemble without positional restraints, with three independent replicas per system.

Trajectories were analysed using GROMACS tools to calculate RMSD, RMSF, radius of gyration and interatomic distances. Protein-ligand interactions and binding affinities were analysed in SeeSAR, and 2D interaction diagrams were generated with the ProteinsPlus PoseView server [56].

### Well-tempered metadynamics simulations

Well-tempered metadynamics simulations were performed with PLUMED 2.9.0 [57], patched to GROMACS 2023 to explore relative protein-ligand motions. A distance-based collective variable was defined as the distance between the centres of mass of the protein and bound ligand. Metadynamics bias was applied using a Gaussian width of 0.1 nm, initial hill height of 0.6 kJ/mol, deposition stride of 200 steps and bias factor of 10. Three independent 100 ns replicas were performed for each system, with collective variable trajectories and deposited bias recoded in COLVAR and HILLS files, respectively.

### **τ**-Random Acceleration Molecular Dynamics (**τ**RAMD) simulations

τRAMD simulations were performed with GROMACS 2020.5-RAMD 2.0 [58], to compare NAD^+^ and AMP unbinding from the AMPK γ1 subunit. A randomly oriented external force was applied to the ligand to accelerate egress from the binding pocket without imposing a predefined dissociation pathway. The receptor was defined as the protein and the ligand as NAD^+^ or AMP. A force of 10 kcal/mol/Å was applied, with force direction evaluated every 50 simulation steps and reassigned if ligand displacement was below 0.0025 nm. Simulations were terminated once the ligand moved more than 6.0 nm from the binding site, and this time was recoded as the dissociation time. For each ligand and binding pose, 30 independent replicas were generated from each starting structure. Dissociation times were analysed to generate cumulative dissociation profiles, residence-time distributions and average residence times for comparison between ligands and binding states.

### Electrostatic mapping

The crystal structure of human AMPKγ1 was obtained from the Protein Data Bank (PDB: 4CFF). Electrostatic surface potential was calculated using the Poisson-Boltzmann method, implemented within the APBS PyMOL plugin [59,60]. Hydrogen atoms, charges and atomic radii were assigned using default APBS/PDB2PQR settings. Electrostatic potential was mapped onto the molecular surface in PyMOL for visualisation.

### Protein sequence alignment

Protein sequences of the AMPK γ1subunit (*PRKAG1* and *SNF4*) were obtained from UniProtKB. Reviewed Swiss-Prot entries were used where available, while unreviewed entries or transcript-derived translations were used for *M. mulatta, G. gallus*, *D. rerio*, *D. melanogaster* and *C. elegans*. Multiple sequence alignment was carried out using Jalview (v 2.11.5.0, University of Dundee), using MAFFT with the L-INS-I strategy. Conservation is indicated by depth of colour from light-to-dark blue.

### Dianthus Spectral Shift Assay

Recombinant AMPKα2β2γ3 complex was expressed in Sf9 insect cells and purified to homogeneity by Ni-affinity and gel-filtration chromatography. AMPK-α2-KD and AurA_KD proteins were expressed in *Escherichia coli* BL21 cells and purified independently. All proteins contained an N-terminal 6×His tag to enable fluorescent labelling with RED-tris-NTA 2nd Generation dye. Labelled proteins (12.5 nM final concentration) were incubated with a 1:1 serial dilution series of NAD or NADH in 384-well microplates. Spectral shift measurements were acquired by determining the fluorescence emission ratio at 670 nm/650 nm over a ligand concentration range of 10 mM to 305 nM. Data were analysed and plotted using GraphPad Prism employing a non-linear regression fit with a one-site total binding model.

## Statistical analyses

Data from at least three independent biological replicates are presented as mean ± s.e.m. and analysed using Welch’s t-test, one-way ANOVA, or other statistical tests as indicated in the figure legends. Multiple comparisons were corrected using Tukey’s post hoc test where appropriate. Statistical analyses were performed using GraphPad Prism 10 software. For enzyme dose-response curves, IC50 values were determined by nonlinear regression (absolute IC50, X = inhibitor concentration) using GraphPad Prism. P < 0.05 was considered statistically significant. Significance is indicated as follows: * *P* < 0.05; ** *P* < 0.01; *** *P* < 0.001; **** *P* < 0.0001; no annotation, not significant.

---

**Figure S1.**
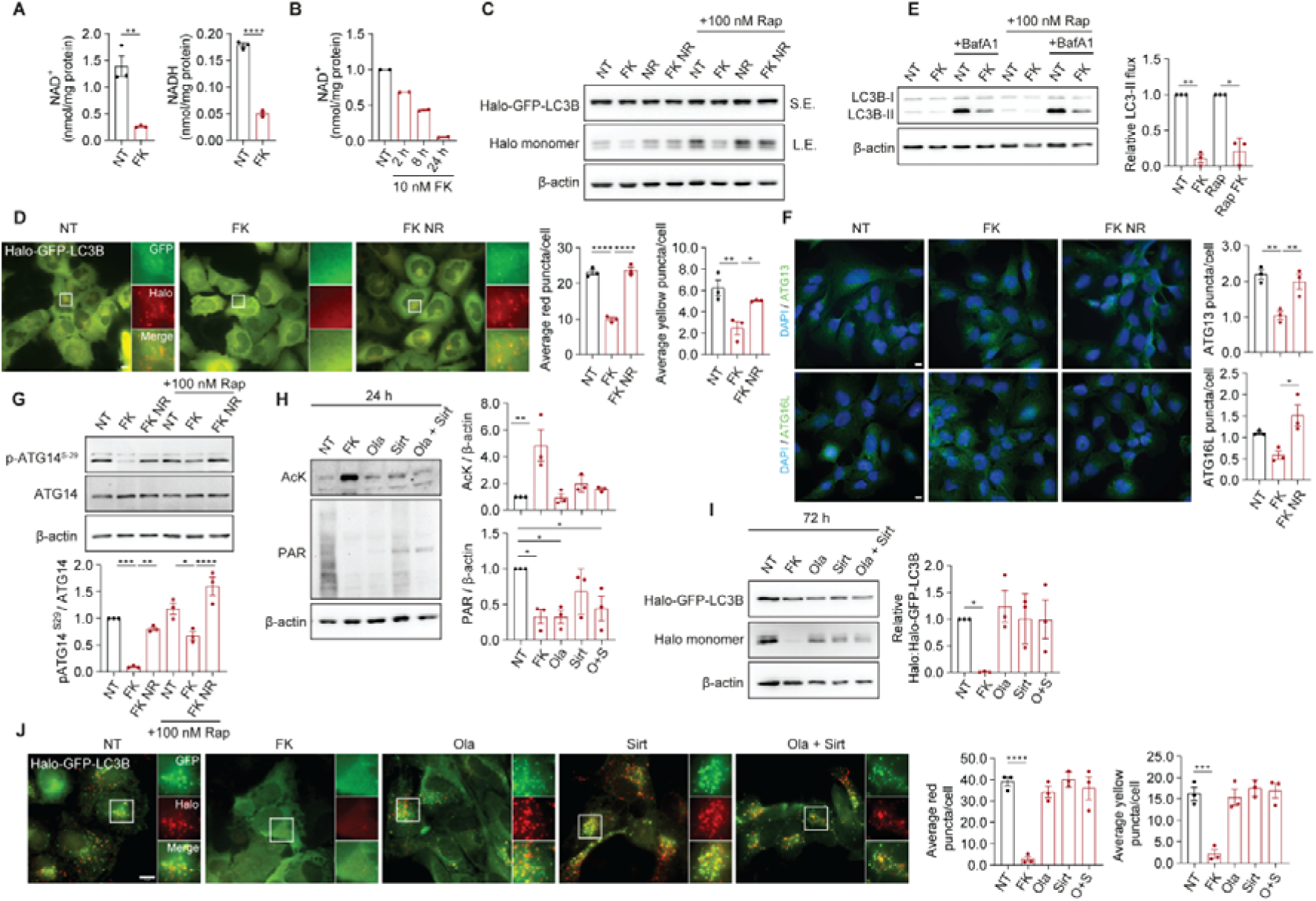
NAD depletion inhibits autophagy, independently of NAD-consuming enzyme activity. **A, B,** Measurement of NAD^+^ and NADH in HMC3 cells treated with 10 nM FK866 (FK) for 24 h (A), and NAD^+^ levels following FK treatment for the indicated timepoints, relative to untreated (B). **C, D,** Immunoblot (C) and immunofluorescence images/quantifications (D) of halo processing assay in HMC3 cells stably expressing Halo-GFP-LC3B. Cells were treated with 10 nM FK for 24 h and/or 1 mM nicotinamide riboside (NR) for 24 h. 100 nM rapamycin (Rap) was added for the final 6 h where indicated. **E,** Immunoblot analysis of autophagy flux in HMC3 cells treated as in C-D with 100 nM Rap and/or 250 nM Bafilomycin A1 (BafA1) where indicated for the final 6 h. **F,** Fluorescence microscopy images and puncta quantification of fixed HMC3 cells labelled with an anti-ATG13 or anti-ATG16L antibody. Cells were treated as in C-D in the presence of 100 nM Rap. **G,** Representative immunoblot and quantification of pATG14Ser-29 in HMC3 cells treated as in C-D. Cells were treated as in C-D. **H,** Representative immunoblot of acetylated lysine (AcK) and Poly-ADP-ribose (PAR) from ARPE-19 cells treated with 100 nM FK, 5 μM Olaparib (Ola) and/or 40 μM Sirtinol (Sirt) for 24 h. **I, J,** Representative immunoblot and quantification (I) or immunofluorescence images/quantification (J) of Halo processing assay in ARPE-19 cells stably expressing Halo-GFP-LC3B, treated as in H for 72 h. Data are mean ± SEM of three independent experiments (A, D-J), data of two independent experiments (B) or immunoblots of a single experiment (C). *P* values were calculated by Welch’s t test (A, E), or ordinary one-way ANOVA and Tukey’s multiple comparisons test (D, F-J). \**P*<0.05; \*\**P*<0.01; \*\*\**P*<0.001; \*\*\*\**P*<0.0001. Scale bars: 10 µm (D, F, J).

**Figure S2.**
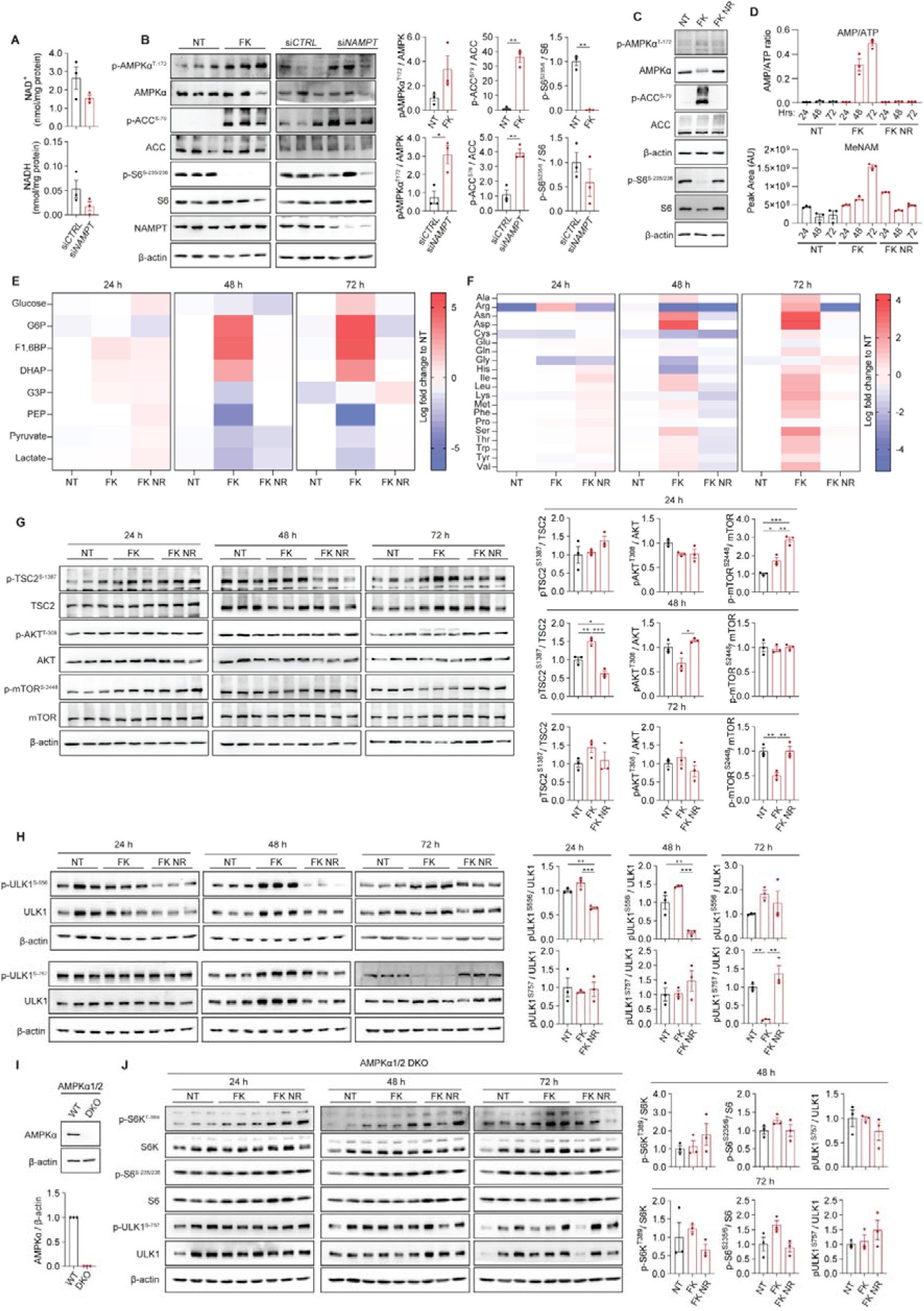
AMPK is required for mTOR suppression by NAD depletion. **A,** Measurement of NAD^+^ and NADH in ARPE-19 cells transfected with *NAMPT* or non-targeting control siRNA. **B,** Representative immunoblots and quantification of indicated proteins from ARPE-19 cells treated with 100 nM FK866 (FK) for 72 h, or transfection with either *NAMPT* or non-targeting control siRNA. **C,** Immunoblot of indicated proteins from HMC3 cells treated with 10 nM FK in the presence or absence of 1 mM nicotinamide riboside (NR) for 24 h. **D,** Metabolite analysis in ARPE-19 cells by LC-MS. Cells were treated with 100 nM FK in the presence of absence of 1 mM nicotinamide riboside NR for the indicated timepoints. Lysates were normalised to 2 × 10L cells/mL. Data show quantified ratio of AMP/ATP peak areas (AU), and the peak area of Methyl-nicotinamide. **E-F,** Heatmaps of glycolysis intermediates (E) and amino acids (F) from ARPE-19 cells by LC-MS, treated as in D. Heatmaps show metabolite log2 fold change relative to untreated (NT) for each timepoint. **G, H,** Immunoblots and quantifications of indicated mTOR-related proteins (G) or ULK1 phosphorylation status (H, S556 or S757) from ARPE-19 cells treated as in D. Immunoblots in H are derived from the same membranes used in Figure 2B and therefore share the same loading control. **I,** Representative immunoblot and quantification of AMPKα from ARPE-19 AMPKα1/α2 WT and DKO cells. **J,** Immunoblots and quantifications of indicated mTOR substrate proteins from ARPE-19 AMPKα1/α2 WT and DKO cells treated as in D. Data are mean ± SEM of three independent experiments (A-B, D, G-J), summary data of three independent experiments (E-F) or immunoblots of a single experiment (C). \**P*<0.05; \*\**P*<0.01; \*\*\**P*<0.001. Immunoblots in G, H and J represent three independent biological replicates per treatment condition (n = 3 per condition, 9 samples total). *P* values were calculated by Welch’s t test (A-B), or ordinary one-way ANOVA and Tukey’s multiple comparisons test (G, H, J).

**Figure S3.**
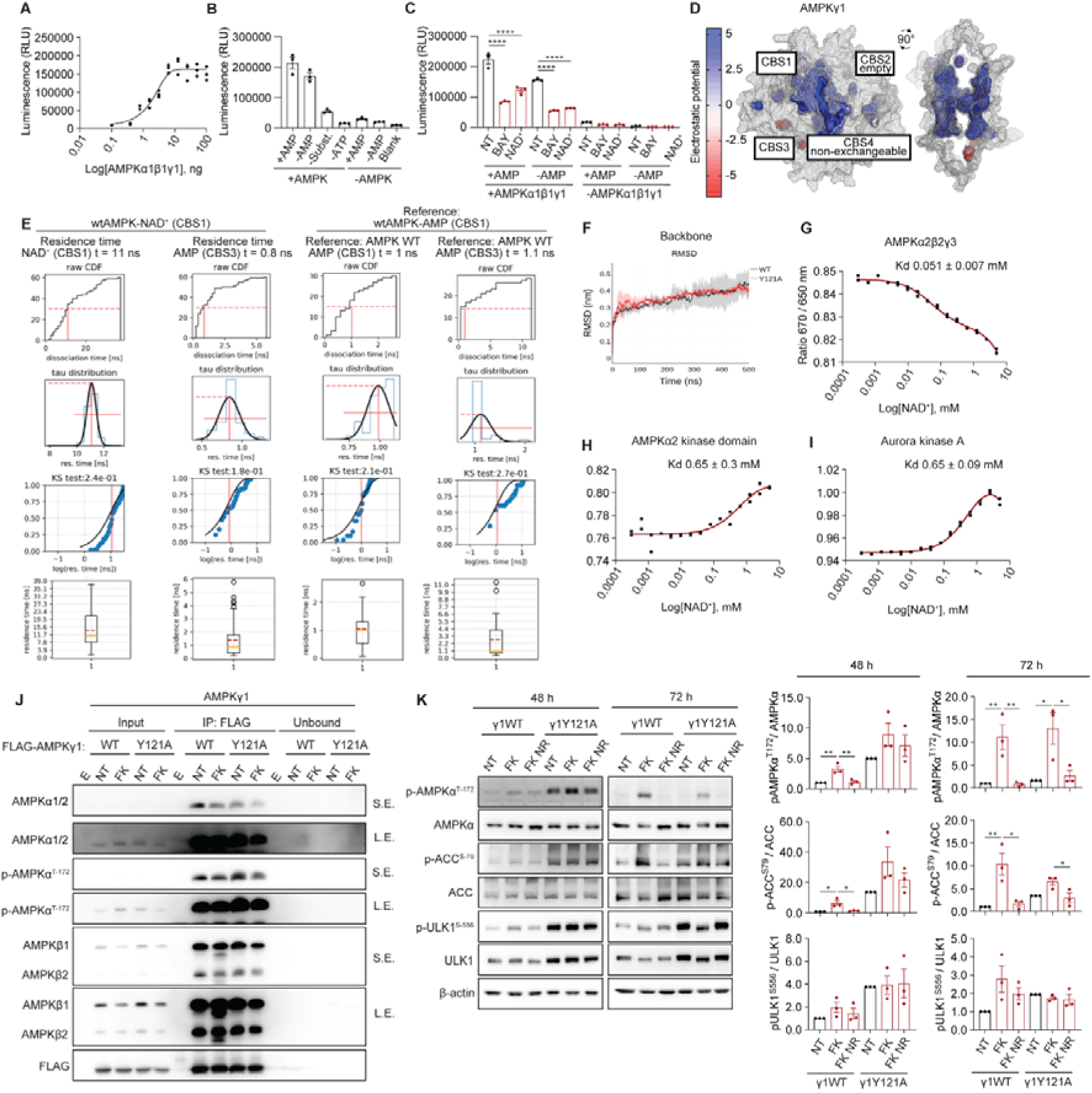
AMPK**γ**1 Y121A mutation desensitises AMPK complexes to NAD depletion. **A-C,** Kinase activity of recombinant human AMPKα1β1γ1 measured using the ADP-Glo^TM^ assay. Activity was measured across increasing enzyme concentrations (0.1-100 ng/well) in the presence of 100 μM AMP (A), with control reactions used to determine background signal and assay performance (B). Kinase activity under no treatment (NT), 350 μM NAD^+^, or 100 nM BAY-3827 was measured in the presence or absence of 100 μM AMP (C). AMPK (10 ng/well) or reaction buffer (blank) was pre-incubated with NAD^+^, BAY-3827, and/or AMP for 1 h before reaction initiation. **D,** Structural model of human AMPKγ1 (PDB: 4CFF). Electrostatic surface potential was calculated using the Poisson-Boltzmann method with the APBS PyMOL. The external surface is shown as a grey mesh, with internal cavities rendered according to electrostatic potential. The positions and binding preferences of the four CBS sites are indicated by annotated boxes. **E,** Calculated residence times of NAD^+^ bound at CBS1 compared with AMP bound at CBS1 or CBS3, determined by τ-random acceleration molecular dynamics (τRAMD). For each ligand/site combination, 30 independent trajectories were analysed. Panels show cumulative dissociation-time distributions, fitted residence-time (τ) distributions, Kolmogorov-Smirnov goodness-of-fit analyses, and summary box plots of dissociation times. NAD^+^ at CBS1 showed a longer predicted residence time than AMP at CBS1 or CBS3, consistent with prolonged engagement at the CBS1 site. **F,** Average root-mean-square-deviation (RMSD) of the AMPKγ1 backbone structure in wild-type (WT, black) and Y121A mutant (red) during 500 ns equilibrium all-atom MD simulations. Data represent averages of three independent replicas; shaded areas indicate SD. **G-I,** Representative binding curves showing interaction of NAD^+^ with recombinant AMPKα2β2γ3 (G), isolated AMPKα2 kinase domain (H), and control Aurora A kinase (I). Kd values were determined using a 1:1 binding model. Note that the direction for the spectral shift change varies depending on the protein and local environment of the labelled fluorophore. **J,** Representative immunoblot of anti-FLAG immunoprecipitates from ARPE-19 cells stably overexpressing WT or Y121A FLAG-AMPKγ1. Cells were treated with no treatment (NT) or 100 nM FK866 for 24 h prior to lysis. Immunoprecipitates were probed for indicated proteins and phosho-markers. E denotes empty (FLAG-beads only) control. **K,** Representative immunoblots and quantifications of indicated proteins from ARPE-19 cells stably expressing WT or Y121A mutant FLAG-AMPKγ1, treated with 100 nM FK866 (FK) in the presence or absence of 1 mM nicotinamide riboside (NR) for 48 or 72 h. The NT baseline of Y121A was scaled to the average NT of WT to account for its higher activity. Data are mean ± SEM of three independent experiments (K), three (A-C) or two (G-I) technical replicates, or immunoblots of a single experiment (J). *P* values were calculated by ordinary one-way ANOVA and Tukey’s multiple comparisons test (C, K). \**P*<0.05; \*\**P*<0.01; \*\*\**P*<0.001; \*\*\*\**P*<0.0001.

**Figure S4.**
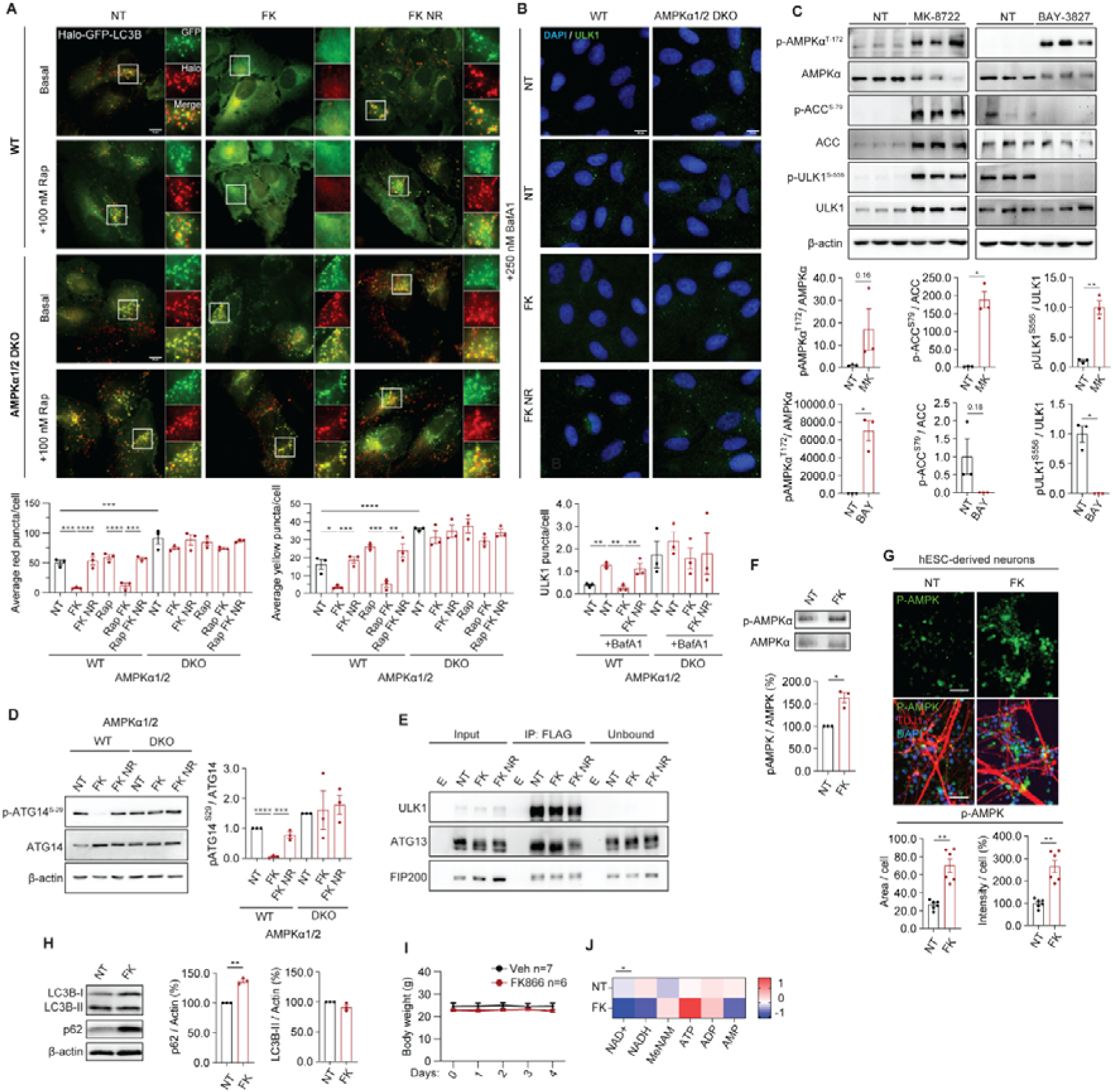
AMPK activation compromises autophagy initiation upon NAD depletion. **A,** Fluorescence microscopy images and quantification of autophagy events (red) and autophagosomes (yellow) in ARPE-19 AMPKα1/α2 WT and DKO cells stably expressing Halo-GFP-LC3B. Cells were treated with 100 nM FK866 (FK) in the presence or absence of 1 mM nicotinamide riboside (NR) for 72 h. For the final 6 h of treatment, 100 nM rapamycin (Rap) was added where indicated. **B,** Fluorescence microscopy images and puncta quantification of fixed ARPE-19 AMPKα1/α2 WT and DKO cells labelled with an anti-ULK1 antibody. Cells were treated as in A, with 250 nM Bafilomycin A1 (BafA1) added for the final 6 h of treatment. **C,** Representative immunoblot and quantification of AMPK-related proteins from ARPE-19 cells treated with 10 μM MK-8722 (MK; AMPK activator) or 5 μM BAY-3827 (BAY; AMPK inhibitor) for 72 h. **D,** Representative immunoblot and quantification of pATG14Ser-29 in ARPE-19 AMPKα1/α2 WT and DKO cells treated as in A. **E,** Immunoblot of indicated proteins from ARPE-19 ULK1 KO cells re-expressing FLAG-ULK1 WT, following immunoprecipitation with anti-FLAG beads. Cells were treated as in A. E = Empty, anti-FLAG beads only. **F-H,** Representative immunoblot (G), immunofluorescence image (H) and quantification (G, H) of p-AMPKαThr172, and representative immunoblot and quantification of LC3B and p62 (I) in hESC-derived neurons, treated with DMSO (NT) or 10 nM FK for the last 6 days of the 4-weeks neuronal differentiation period. **I-J,** Body weight (I) and heatmaps of indicated metabolites (log2 relative to NT, J) of mice treated with vehicle (NT) or 30 mg/kg FK866 once daily for 5 days. Data are mean ± SEM of three independent experiments (A-D, F-H), mean ± SD of n=13 (I, 7 control, 6 treated), summary data of n=10 (J, 5 control, 5 treated), or representative immunoblot of a single experiment (E). *P* values were calculated by ordinary one-way ANOVA and Tukey’s multiple comparisons test (A-B, D), Welch’s t-test (C, F-H), two-way ANOVA (I), or unpaired two-tailed t-test with Welch’s correction on log-transformed data with a two-stage linear step-up method (J). \**P*<0.05; \*\**P*<0.01; \*\*\**P*<0.001; \*\*\*\**P*<0.0001. Scale bars: 10 µm (A, B) or 50 µm (G).

